# Novel genetic modules encoding high-level antibiotic-free protein expression in probiotic lactobacilli

**DOI:** 10.1101/2022.08.04.502766

**Authors:** Sourik Dey, Marc Blanch-Asensio, Sanjana Balaji Kuttae, Shrikrishnan Sankaran

## Abstract

Lactobacilli are ubiquitous in nature, often beneficially associated with animals as commensals and probiotics, and are extensively used in food fermentation. Due to this close-knit association, there is considerable interest to engineer them for healthcare applications in both humans and animals, for which high-performance and versatile genetic parts are greatly desired. For the first time, we describe two genetic modules in *Lactiplantibacillus plantarum* that achieve high-level gene expression using plasmids that can be retained without antibiotics, bacteriocins or genomic manipulations. These include (i) a promoter, P_*tlpA*_, from a phylogenetically distant bacterium, *Salmonella typhimurium*, that drives up to 5-fold higher level of gene expression compared to previously reported promoters and (ii) multiple toxin-antitoxin systems as a self-contained and easy-to-implement plasmid retention strategy that facilitates the engineering of tunable transient Genetically Modified Organisms. These modules and the fundamental factors underlying their functionality that are described in this work will greatly contribute to expanding the genetic programmability of lactobacilli for healthcare applications.

## INTRODUCTION

Lactobacilli are gram-positive rod-shaped lactic acid bacteria (LAB), typically found in humans and animals as commensals. Their stress tolerant phenotypic traits allow them to colonize a wide range of host microenvironments, like the gut, skin, vagina, nasal and oropharyngeal cavity (Ma *et al*., 2012; Turroni *et al*., 2014) often providing health benefits in the form of anti-inflammatory, anti-pathogenic and immunomodulatory activities (Darby and Jones, 2017; Bibalan *et al*., 2017). Due to this, they are one of the largest classes of probiotics and several species are being clinically tested for treating a variety of diseases like ulcerative colitis (Zocco *et al*., 2006), mastitis(Jiménez *et al*., 2008), atopic dermatitis (Rosenfeldt *et al*., 2003), bacterial vaginosis (Mastromarino *et al*., 2009) and periodontitis (Teughels *et al*., 2013). Apart from their health benefits, lactobacilli are also vital for numerous fermentation processes in the food industry, for example in the production of yogurt (Ashraf and Shah, 2011), cheese (Kasımoğlu *et al*., 2004), sourdough bread (Plessas *et al*., 2008), beer (Chan *et al*., 2019) and wine (du Toit *et al*., 2011). Due to this ubiquity in our lives, there is considerable interest to genetically enhance and expand the capabilities of these bacteria for healthcare applications (Pedrolli *et al*., 2019). For instance, lactobacilli are being engineered as live biotherapeutic products (LBPs) that produce and deliver drugs right at the site of diseases like ulcerative colitis (de Vos, 2011), Human Immunodeficiency Virus (HIV) infection (Watterlot *et al*., 2010) and respiratory infections (Janahi *et al*., 2018). They are also prominent candidates for the development of mucosal vaccines in which they are engineered to either display heterologous antigens on their surface or to secrete them (LeCureux and Dean, 2018). These food-grade *Lactobacillus* vaccine vectors would be cheap to produce and can be easily administered orally or intranasally, improving the ability to deploy them both in humans and animals. Examples of infectious diseases against which such vaccines are under development include anthrax, infantile diarrhea, pneumonia and viral infections like, HIV, HPV, influenza and coronavirus (LeCureux and Dean, 2018; Wang *et al*., 2020). Finally, to track these therapeutic bacteria within the body and study their colonization and clearance profiles, there is considerable interest to make them express reporter proteins that can be imaged *in situ* (Landete *et al*., 2015; Salomé-Desnoulez *et al*., 2021).

Despite such potential, the main limitations for engineering lactobacilli are the scarcity of well-characterized genetic parts and insufficient understanding of biochemical pathways required to build the type of genetic circuits that have been demonstrated in *E. coli* (Elowitz and Leibler, 2000; Wang *et al*., 2011) and *B. subtilis* (Courbet *et al*., 2015; Castillo-Hair *et al*., 2019). Over two decades of painstaking investigation and screening across phylogenetically close bacteria have generated a handful of reliable parts for use in lactobacilli such as constitutive and inducible promoters, operators, replicons, retention-modules, signal peptides etc. Most of these have been developed in a few species that were found to be amenable to genetic modification, among which *Lactiplantibacillus plantarum* (Zheng *et al*., 2020) is widely reported (Siezen and van Hylckama Vlieg, 2011). While genomic integration of genes has been demonstrated in these bacteria, the greatest versatility of functions has been achieved using plasmids. Excellent progress has been made in establishing plasmid backbones with low, medium and high copy number replicons (Tauer *et al*., 2014), constitutive promoters with a wide range of expression strengths (Rud *et al*., 2006), a few inducible promoters that can be triggered by peptides (Halbmayr *et al*., 2008) or sugars (Heiss *et al*., 2016), signal peptides sequences enabling protein secretion (Mathiesen *et al*., 2009) or surface display (Mathiesen *et al*., 2020) and food-grade plasmid retention systems based on resistance to external stressors (e.g. bacteriocins) (Takala and Saris, 2002; Allison and Klaenhammer, 1996) or auxotrophy complementation requiring genomic knockout of a metabolic gene and providing it in the plasmid (Nguyen *et al*., 2011; Chen *et al*., 2018). However, the available set of well-characterized genetic parts is still minuscule compared to the toolbox of *E. coli* and needs to be expanded in order to improve the performance and versatility of *Lactobacillus* engineering for healthcare applications.

In this work, we introduce 2 new versatile and powerful genetic parts to expand the capabilities of *Lactobacillus* engineering - (i) a novel constitutive promoter from a phylogenetically distant *Salmonella* species that drive protein expression at levels considerably higher than previously reported strong *L. plantarum* promoters and (ii) toxin-antitoxin systems as an alternative strategy for plasmid retention that does not require manipulating the bacterial genome. Unique features of the novel promoter sequence are discussed, which can lead to new design criteria for improving promoter strengths in lactobacilli. The toxin-antitoxin systems introduce a thus-far unexplored modality of plasmid retention in lactobacilli that enables the generation of temporary Genetically Engineered Microorganisms (GEMs), desirable for medical and food-grade applications. These parts and the fundamental insights gained in their characterization will strongly aid in expanding the genetic programmability of lactobacilli.

## MATERIALS AND METHODS

### Strain, Media and Plasmids

*L. plantarum* WCFS1 was used as the parent strain for promoter strength and plasmid retention characterization. The strain was maintained in the De Man, Rogosa and Sharpe (MRS) media. The culture media, antibiotics and complementary reagents were purchased from Carl Roth Gmbh, Germany. Growth media was supplemented with 10 μg/mL of erythromycin to culture engineered *L. plantarum* WCFS1 strains. The plasmids pSIP403 and pLp_3050sNuc used in this study were a kind gift from Prof. Lars Axelsson (Addgene plasmid # 122028) (Sørvig *et al*., 2005a) and Prof. Geir Mathiesen (Addgene plasmid # 122030) (Mathiesen *et al*., 2009) respectively. The plasmid pTlpA39-Wasabi was a kind gift from Prof. Mikhail Shapiro (Addgene plasmid # 86116) (Piraner *et al*., 2017). The plasmid pUC-GFP-AT was a kind gift from Prof. Chris Barnes (Addgene plasmid # 133306) (Fedorec *et al*., 2019). The sequence verified genetic constructs created in this study have been maintained in *E. coli* DH5α.

### Molecular Biology

The genetic constructs developed in this study are based on the pLp3050sNuc/ pSIP403 vector backbone. The HiFi Assembly Master Mix, Quick Blunting Kit and the T4 DNA Ligase enyzme were purchased from New England BioLabs (NEB, Germany). PCR was performed using Q5 High Fidelity 2X Master Mix (NEB) with primers purchased from Integrated DNA Technologies (IDT) (Leuven, Belgium). Oligonucleotide gene fragments were purchased as eBlocks from IDT (Coralville, USA). These were codon optimized for maximal expression in the host strain using the IDT Codon Optimization Tool (Coralville, USA). Plasmid extraction and DNA purification were performed using kits purchased from Qiagen GmbH (Hilden, Germany) and Promega GmbH (Walldorf, Germany) respectively. The general schematic of plasmid construction for this study has been shown in Supplementary Figure S1. The promoter sequences used in this study are provided in Supplementary Table S1 and the nucleotide sequences of the toxin-antitoxin modules have been highlighted in Supplementary Table S2.

### *L. plantarum* WCFS1 Competent Cell Preparation and DNA Transformation

A single colony of *L. plantarum* WCFS1 was inoculated in 5 mL of MRS media and cultured overnight at 37 °C with shaking (250 rpm). The primary culture was diluted in a 1:50 (v/v) ratio in a 25 mL secondary culture composed of MRS media and 1% (w/v) glycine premixed in a 4:1 ratio. The secondary culture was incubated at 37 °C, 250 rpm until OD_600_ reached 0.8, following which the cells were pelleted down by centrifuging at 4000 rpm (3363 × g) for 10 min at 4°C. The pellet was washed twice with 5 mL of ice-cold 10 mM MgCl_2_ and then washed twice with 5 mL and 1 mL of ice-cold Sac/Gly solution [10% (v/v) glycerol and 1 M sucrose mixed in a 1:1 (v/v) ratio] respectively. Finally, the residual supernatant was discarded, and the pellet resuspended in 500 μL of Sac/Gly solution. The competent cells were then dispensed in 60 μL aliquots for DNA transformation. For all transformations, 1 μg of dsDNA were added to the competent cells and then transferred to chilled 2 mm gap electroporation cuvettes (Bio-Rad Laboratories GmbH, Germany). Electroporation transformation was done with a single pulse at 1.8 kV, after which 1 mL of lukewarm MRS media was immediately added. The mixture was kept for incubation at 37 °C, 250 rpm for a recovery period of 3 h. Following the recovery phase, the cells were centrifuged at 4000 rpm (3363 × g) for 5 min, 800 μL of the supernatant discarded, and 200 μL of the resuspended pellet was plated on MRS Agar supplemented with 10 μg/mL of Erythromycin. The plates were incubated at 37 °C for 48 h to allow the growth of distinct single colonies.

### Direct cloning in *L. plantarum* WCFS1

To obtain sufficient plasmid quantities (~1 μg) for transformation in *L. plantarum* WCFS1, a modified direct cloning method (Spath *et al*., 2012) involving PCR-based amplification and circularization of recombinant plasmids was used. Plasmids were constructed and transformed directly in *L. plantarum* WCFS1 strain using a DNA assembly method. Complementary overhangs for HiFi Assembly were either created using PCR primers or synthesized as custom designed eBlocks. Purified overlapping DNA fragments were mixed with the HiFi DNA Assembly Master Mix and assembled as recommended in the standard reaction protocol from the manufacturer. The assembled DNA product was then exponentially amplified by another round of PCR using a pair of primers annealing specifically to the insert segment. 5 μl of the HiFi assembly reaction was used as a template for this PCR amplification of the assembled product (100 μl final volume). The purified PCR product was then subjected to phosphorylation using the Quick Blunting Kit. 2000 ng of the purified PCR product was mixed with 2.5 μl of 10X Quick blunting buffer and 1 μl of Enzyme Mix (Milli-Q water was added up to 25 μl). The reaction was incubated first at 25 °C for 30 minutes and then at 70 °C for 10 minutes for enzyme inactivation. Next, phosphorylated products were ligated using the T4 ligase enzyme. 6 μl of the phosphorylated DNA was mixed with 2.5 μl of 10X T4 Ligase Buffer and 1.5 μl of T4 Ligase enzyme (Milli-Q water was added up to 25 μl). Two ligation reactions were performed per cloning (25 μl each). The respective reactions were incubated at 25 °C for 2 hours and then at 70 °C for 30 min for enzyme inactivation. The ligated reactions were mixed together and purified. In order to concentrate the final purified product, three elution rounds were performed instead of one. Each elution was based on 10 μl of Milli-Q water. The concentration of the ligated purified product was measured using the NanoDrop Microvolume UV-Vis Spectrophotometer (ThermoFisher Scientific GmbH, Germany). Finally, 1000 ng of the ligated product were transformed into *L. plantarum* WCFS1 electrocompetent cells, resulting in a transformation efficiency of 2 – 3 × 10^2^ cfu/μg.

Notably, since *L. plantarum* harbors 3 endogenous plasmids (Van Kranenburg *et al*., 2005), sequencing was performed on PCR amplified sections. In detail, colonies of interest were inoculated in MRS supplemented with 10 μg/mL of Erythromycin and grown overnight at 37 °C. The following day, 1 mL of the culture was pelleted down, and the supernatant was discarded. Next, a tip was used to collect a tiny part of the pellet, which was used as a template for PCR (100 μL final volume). Finally, PCR products were purified and sent for Sanger sequencing to Eurofins Genomics GmbH (Ebersberg, Germany) by opting for the additional DNA purification step.

### Microplate reader Setup for Thermal Gradient Analysis

Bacterial cultures were cultivated in 5 mL of MRS media (supplemented with 10 μg/mL erythromycin) at 30°C with continuous shaking (250 rpm). The following day, cultures were diluted to 0.1 OD_600_ in 3 mL of antibiotic supplemented fresh MRS media and propagated at 30°C, 250 rpm. At OD_600_ = 0.3, the cultures were dispensed into Fisherbrand™ 0.2mL PCR Tube Strips with Flat Caps (Thermo Electron LED GmbH, Germany) and placed in the Biometra Thermocycler (Analytik Jena. GmbH, Germany). For the P_spp_-mCherry construct, 25 ng/mL of the 19 amino acid Sakacin P inducer peptide (SppIp) with the sequence NH_2_-MAGNSSNFIHKIKQIFTHR-COOH (GeneCust, France) was added to the culture and thoroughly vortexed before preparing the aliquots. The thermal assay was set at a temperature gradient from 31°C to 41°C with regular increment of 2°C. The lid temperature was set at 50°C to prevent the evaporation of the liquid and maintain a homogeneous temperature in the spatially allocated PCR tubes. After a time interval of 18 h, the PCR strips were centrifuged in a tabletop minicentrifuge (Biozym GmbH, Germany) to pellet down the cells and discard the supernatant. The cells were then resuspended in 200 μL of 1X PBS and added to the clear bottom 96-well microtiter plate (Corning® 96 well clear bottom black plate, USA). The samples were then analyzed in the Microplate Reader Infinite 200 Pro (Tecan Deutschland GmbH, Germany) and both the absorbance (600 nm wavelength) and mCherry fluorescence intensity (Ex_λ_ / Em_λ_ = 587 nm/625 nm) were measured. The z-position and gain settings for recording the mCherry fluorescent intensity were set to 19442 μm and 136 respectively. Fluorescence values were normalized with the optical density of the bacterial cells to calculate the Relative Fluorescence Units (RFU) using the formula RFU = Fluorescence/OD_600_.

### Fluorescence Microscopy Analysis

Bacterial cultures were grown overnight in 5 mL of MRS media (supplemented with 10 μg/mL erythromycin) at 37°C with continuous shaking (250 rpm). The following day, the OD_600_ of the P_spp_-mCherry construct was measured and subcultured at OD_600_ = 0.01. When the P_spp_-mCherry bacterial culture reached OD_600_ =0.3, it was induced with 25 ng/mL of SppIp and the remaining constructs were subcultured in fresh media at 0.01 OD_600_. All the cultures were then allowed to grow for 18 h under the same growth conditions (37°C, 250 rpm) to prevent any heterogeneity in promoter strength expression due to differential growth parameters. Later, 1 mL of the cultures were harvested by centrifugation (15700 × g, 5 min, 4 °C), washed twice with Dulbecco’s 1X PBS (Phosphate Buffer Saline) and finally resuspended in 1 mL of 1X PBS. 10 μL of the suspensions were placed on glass slides of 1.5 mm thickness (Paul Marienfeld GmbH, Germany) and 1.5H glass coverslips (Carl Roth GmbH, Germany) were placed on top of it. The samples were then observed under the Plan Apochromat 100X oil immersion lens (BZ-PA100, NA 1.45, WD 0.13 mm) of the Fluorescence Microscope BZ-X800 (Keyence Corporation, Illinois, USA). The mCherry signal were captured in the BZ-X TRITC filter (model OP-87764) at excitation wavelength of 545/25 nm and emission wavelength of 605/70 nm with a dichroic mirror wavelength of 565 nm. The images were adjusted for identical brightness and contrast settings and were processed with the FiJi ImageJ2 software.

### Flow Cytometry Analysis

Quantification of fluorescent protein expression levels of the strains were performed using Guava easyCyte BG flow-cytometer (Luminex, USA). Bacterial cultures subjected to the same treatment conditions mentioned above were used for Flow Cytometry analysis. 1 mL of the bacterial suspensions were harvested by centrifugation at 13000 rpm (15700 × g). The supernatant was discarded and the pellet was resuspended in 1 mL of sterile Dulbecco’s 1X PBS. The samples were then serially diluted by a 10^4^ Dilution Factor (DF) and 5,000 bacteria events were recorded for analysis. Experiments were performed in triplicates on three different days. During each analysis, the non-fluorescent strain carrying the empty vector was kept as the negative control. A predesigned gate based on forward side scatter (FSC) and side scatter (SSC) thresholding was used to remove debris and doublets during event collection and analysis. mCherry fluorescence intensity was measured using excitation by a green laser at 532 nm (100 mW) and the Orange-G detection channel 620/52 nm filter was used for signal analysis. The gain settings used for the data recording were, Forward Scatter (FSC) – 11.8; Side Scatter (SSC) −4, and Orange-G Fluorescence – 1.68. The compensation control for fluorescence recording was set at 0.01 with an acquisition rate of 5 decades. Data analysis and representation were done using the Luminex GuavaSoft 4.0 software for EasyCyte.

### Toxin/Antitoxin Module based Plasmid Construction

Similar to previous reports in *E. coli* (Fedorec *et al*., 2019), the effect of Txe/Axe (toxin/antitoxin) module from *E. faecium* (Grady and Hayes, 2003) was tested in *L. plantarum* WCFS1 to test its capability for antibiotic-free plasmid retention. TA Finder version 2.0 tool (Xie *et al*., 2018) was used to select further type-II TA (Toxin/Antitoxin) systems present in *Lactobacillus* genomes. *L. acidophilus*, *L. crispatus*, *L. casei*, *L. reuteri*, and *L. plantarum* WCFS1 genomes were retrieved from NCBI Genome. TA systems harbored within these genomes were mined using the default parameters of TA Finder. Only TA systems annotated by NCBI BlastP were selected as test candidates. The TA systems YafQ/DinJ, HicA/HicB, HigB/HigA, MazF/MazE from *L. casei, L. acidophilus and L. plantarum* WCFS1 were selected for further testing and analysis.

Txe/Axe system was amplified by PCR from the plasmid pUC-GFP-AT (Fedorec *et al*., 2019). DinJ/YafQ and HicA/HicB systems were synthesized as custom-designed eBlocks. HigA/HigB and MazE/MazF were amplified from the genome of *L. plantarum* WCFS1. TA systems were inserted into the P_*tlpA*_-mCherry plasmid, generating the plasmids P_*tlpA*_-mCherry-Txe/Axe, P_*tlpA*_-mCherry-YafQ/DinJ, P_*tlpA*_-mCherry-HicA/HicB, P_*tlpA*_-mCherry-HigB/HigA, P_*tlpA*_-mCherry-MazF/MazE For constructing the combinatorial TA module (P_*tlpA*_-mCherry Combo), the best performing endogenous and non-endogenous TA systems recorded after 100 generations (MazF/MazE and YafQ/DinJ) were subcloned and integrated into the same plasmid in reverse orientations.

### TA Mediated Plasmid Retention Analysis

The TA module containing constructs were inoculated in 5 mL cultures of 10 μg/mL erythromycin supplemented MRS media and incubated overnight at 37°C with continuous shaking (250 rpm). The following day, the constructs were subcultured at an initial OD_600_ = 0.01 in fresh MRS media (both with and without antibiotic supplementation). The bacterial cultures were incubated for 12 consecutive days with a daily growth period of 24 h ensuring an average of ~8 generations per day, until crossing the final threshold of 100 generations. Sample preparation for flow cytometry analysis was conducted according to the protocol mentioned before. The mCherry positive cell population directly correlated to the bacterial population retaining the engineered plasmid. The entire experiment was repeated in biological triplicates.

To cross-check the flow cytometry analysis, the bacterial cultures grown for 100 generations without antibiotic supplementation were centrifuged and resuspended in 1 mL of sterile Dulbecco’s 1X PBS. The resuspended bacterial solutions were diluted (DF=10^6)^ and plated on MRS Agar plates supplemented without antibiotic and incubated in a static incubator for 48 h. The plates were then imaged using the GelDocumentation System Fluorchem Q (Alpha Innotech Biozym Gmbh, Germany) both in the Ethidium Bromide channel (Ex_λ_/Em_λ_ = 300 nm/600 nm) and Cy3 channel (Ex_λ_/Em_λ_ = 554 nm/568 nm) to visualize the cell population producing mCherry fluorescence. The fluorescent bacterial subpopulation on the non-selective MRS agar medium correlated to the plasmid retention frequency of the respective TA systems in the absence of selection pressure.

### Growth Rate Measurements

For studying the influence of the heterologous protein production and toxin-antitoxin modules on the bacterial growth rate, bacterial cultures were cultivated overnight in antibiotic supplemented MRS media at 37°C with continuous shaking (250 rpm). Following day, the bacterial cultures were subcultured in secondary cultures at an initial OD_600_ = 0.01. After 4 h incubation at 37°C, the OD_600_ of the cultures reached 0.1 and 200 μL of the cultures were distributed in UV STAR Flat Bottom 96 well microtiter plates (Greiner BioOne GmbH, Germany). The 96 well assay plate was placed in the Microplate Reader with constant shaking conditions at an incubation temperature of 37°C. The kinetic assay was set to record the absorbance of the bacterial cultures at 600 nm wavelength with an interval of 10 min for an 18 h time duration. The experiment was conducted in triplicates on three independent days.

### Bioinformatic analysis

All genome sequence included in the phylogenetic analysis were retrieved from NCBI Genome. The phylogenetic tree was built using the web server for genome-based prokaryote taxonomy “Type (Strain) Genome Server” (TYGS), restricting the analysis only to the sequences provided (Meier-Kolthoff and Göker, 2019). The Genome BLAST Distance Phylogeny (GBDP) tree, based on 16S rDNA gene sequences, was obtained. The Interactive Tree of Life (iTOL) tool was used for the display, annotation, and management of the phylogenetic tree (Letunic and Bork, 2007).

For the multiple sequence alignment, protein sequences of the σ70 subunits from *L. plantarum*, *E. coli* and *S. typhimurium* RNA polymerases were first retrieved from Uniprot. Sequences were aligned using the tool MUSCLE (Edgar, 2004). Jalview was used to visualize and edit the multiple sequence alignment (Waterhouse *et al*., 2009).

SnapGene was used to identify DNA sequences similar to P_*tlpA*_ within the genome of *L. plantarum* WCFS1 using the feature “Find Similar DNA Sequences”. The search allowed a mismatch or gap/insertion every 4 bases. BPROM, an online tool for predicting bacterial promoters, was used to identify the −35 and −10 boxes within this promoter (Madeira *et al*., 2022). BlastP was used to identify the protein encoded by the gene driven by this promoter. Promoter alignment was performed using MUSCLE (Edgar, 2004).

## RESULTS AND DISCUSSIONS

### *P_PtlpA_* Promoter from Salmonella drives high-level constitutive expression

The strongest promoters in lactobacilli have been found by either screening the genome of the host strain (Rud *et al*., 2006; Bron *et al*., 2004) or adapting those driving high-level protein expression in phylogenetically close lactic acid bacteria (Russo *et al*., 2015) (Figure 1A). In the few reports where promoters from phylogenetically distant species like *P. megaterium* (*P_xylA_*) or *E. coli* (*P_T7_* from lambda phage) (Heiss *et al*., 2016) have been tested, expression levels were found to be comparatively low. Contrary to this trend, we serendipitously stumbled upon a promoter (P*_tlpA_)* from the phylogenetically distant gram-negative *Salmonella typhimurium* (Figure 1A) capable of driving protein expression at levels higher than previously reported strong promoters in *L. plantarum* WCFS1. In Salmonella, P_*tlpA*_ along with its repressor is capable of thermo-responsively regulating gene expression and this functionality had been previously transferred to *E. coli* for therapeutic purposes (Piraner *et al*., 2017; Hurme *et al*., 1997). To test whether the P_*tlpA*_ promoter would be a suitable candidate for driving transcription in *L. plantarum*, a fluorescent reporter protein (mCherry) was cloned downstream of this promoter. The promoter surprisingly seemed to constitutively drive a high-level of protein expression with a mild degree of thermal regulation (<5-fold increase from 31 °C to 39 °C) (Figure 1B). Next the repressor based thermo-responsive functionality was tested in *L. plantarum*, by creating the pTlpA39 plasmid, with the P_*tlpA*_ promoter driving expression of mCherry and the codon optimized TlpA repressor being expressed constitutively by the P_*48*_ promoter (Rud *et al*., 2006). However, the pTlpA39 plasmid showed no significant repression of mCherry at lower temperature gradients in comparison to its repressor-free counterpart (Supplementary Figure S2D). Most remarkably, flow cytometry and fluorescence spectroscopy analysis revealed that mCherry expression levels driven by the P_*tlpA*_ promoter significantly exceeded the levels driven by some of the strongest promoters previously reported in *L. plantarum* - P_*23*_ (Meng *et al*., 2021), P_*48*_ (Rud *et al*., 2006), P_*spp*_ (Sørvig *et al*., 2003) and P_*Tuf*_ (Spangler *et al*., 2019) (Figure 1C, Supplementary S3A). At 31 °C, mCherry expression levels were at least 2-fold higher than these other promoters, while this increased to 5-fold at 39 °C (Figure 1D, Supplementary Figure S3B). All constitutive promoters (P_*23*_, P_*48*_, P_*Tuf*_) were mildly thermo-responsive, while the inducible promoter (*P_spp_*) was not (Supplementary Figure S4A). To check whether such high gene expression can be driven by other phylogenetically distant thermo-responsive promoters, we tested the well-known heat inducible *pR* and *pL* promoters from *E. coli* lambda phage. However, only low levels of mCherry expression were observed with these promoters (Supplementary Figure S2A, Supplementary S2B). Fluorescence spectroscopy revealed that the strength of the P_*tlpA*_ promoter at 37 °C was 26- and 39-fold higher than the *pR* and *pL* promoter, respectively (Supplementary Figure S2C).

**Figure 1.**
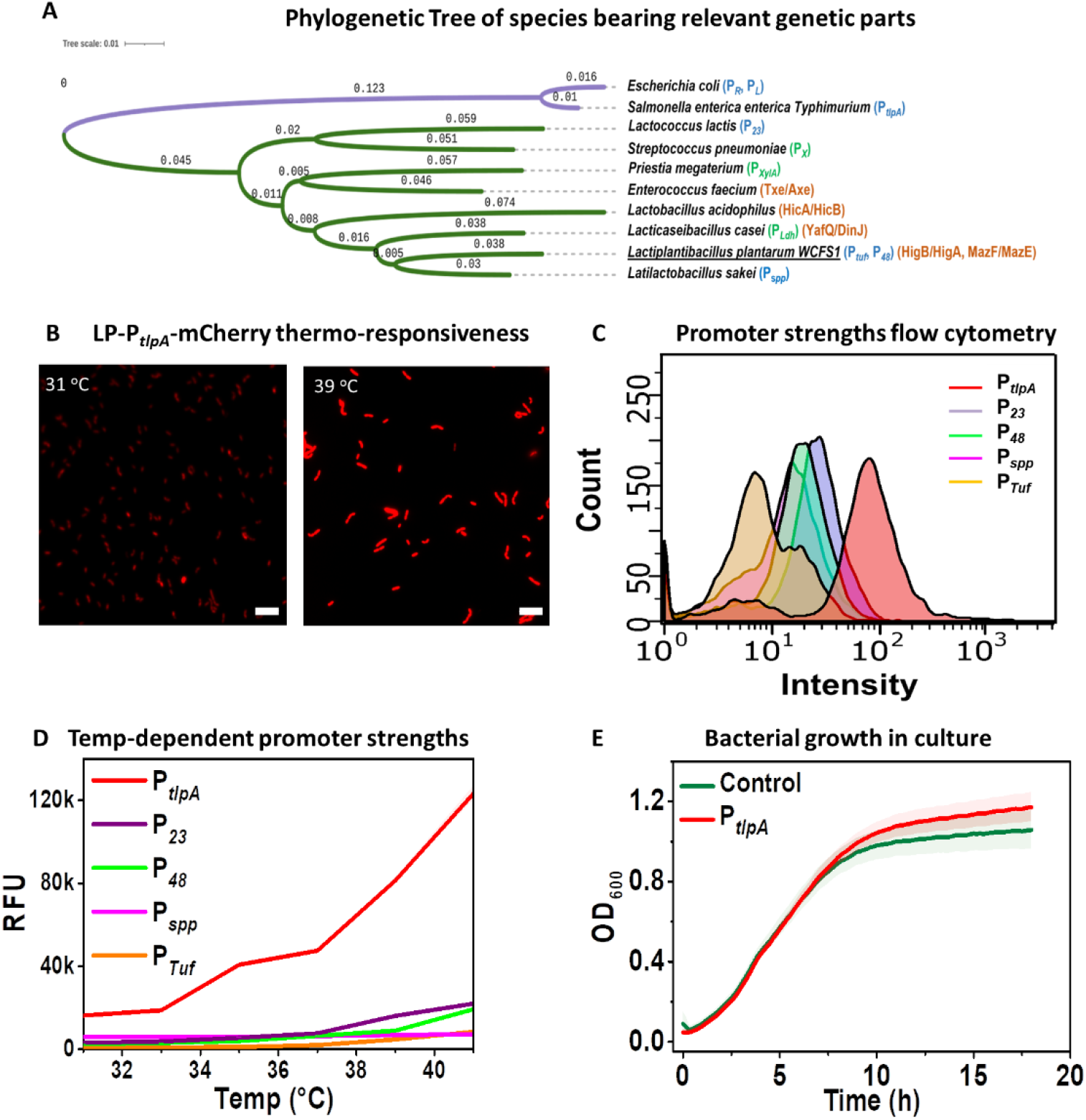
(A) Phylogenetic tree highlighting the distances between species from which various genetic parts have been tested in *L. plantarum*. Purple clade corresponds to Gram-negative bacteria. Green clade corresponds to Gram-positive bacteria. Promoters tested in this study are labelled in blue. Promoters tested by others in *L. plantarum* are labelled in green. Orange labels correspond to the TA systems tested in this study. (B) Fluorescence microscopy of P_*tlpA*_ driven mCherry expression in *L. plantarum* WCFS1 cultivated at 31°C and 39°C for 18 h. Scale bar = 10 μm (C) Flow Cytometry analysis of P_*tlpA*_, P_*23*_, P_*48*_, P_*spp*_ and P_*Tuf*_ driven mCherry expression in *L. plantarum* WCFS1 after 18 h incubation at 37°C. (D) Fluorescence spectroscopy analysis of the P_*tlpA*_, P_*23*_, P_*48*_, P_*spp*_ and P_*Tuf*_ driven mCherry expression after 18 h incubation at temperatures ranging from 31°C to 41°C. (E) Growth rate (OD_600_) measurement of *L. plantarum* WCFS1 strains containing a control plasmid and P_*tlpA*_-mCherry for 18 h at 37°C. In (C) and (D), the solid lines represent mean values, and the lighter bands represents standard deviations calculated from three independent biological replicates.

Another important factor for high-level gene expression driven by *P_tlpA_* is that the spacer length between the ribosome binding site (RBS, 5’-AGGAGA-3’) and the start codon needs to be different in *L. plantarum* compared to *E. coli*. In *E. coli* this spacer length of 6 bp has been previously reported (Piraner *et al*., 2017; Kan et al., 2020; Chee et al., 2022; Rottinghaus et al., 2022), whereas in *L. plantarum* a 9 bp spacer improves expression levels by 25-fold compared to a 6 bp spacer (Supplementary Figure S4B), in accordance with previous reports (Tauer *et al*., 2014). Despite the high level of protein expression driven by P_*tlpA*_ with a 9 bp spacer, the growth rate of this strain at 37 °C was similar to that of the empty vector control strain, suggesting that this protein overexpression did not metabolically overburden the cell (Figure 1E).

To understand why *P_tlpA_* drives gene expression in *L. plantarum*, we looked into its function in Salmonella, where it is a promoter of the σ70 sigma factors. (Dawoud et al., 2017). This family of sigma factors is involved in regulating the expression of housekeeping genes in most prokaryotes, including lactobacilli (Todt *et al*., 2012). Multiple Sequence Alignment (MSA) among the major RNA polymerase σ70 proteins (RpoD) of *E. coli*, S. typhimurium, and *L. plantarum* strains (Figure 2A) revealed significant similarity between the domain-2 and domain-4 regions, responsible for binding to the −10 and −35 regions of the promoter during transcription initiation. Interestingly, Gaida *et al*., (2015) showed that, when expressed in *E. coli*, the *L. plantarum* RpoD can recruit *E. coli*’s RNA polymerase to initiate transcription from a wide variety of heterologous promoters. Compared to sigma factors from six other bacteria, they found that the *L. plantarum* RpoD was the most promiscuous and helped to enlarge the genomic space that can be sampled in *E. coli*. These analyses explain why the P_*tlpA*_ promoter from a phylogenetically distant species functions in *L. plantarum* but does not necessarily reveal how it drives such high expression levels compared to previously reported promoters.

**Figure 2.**
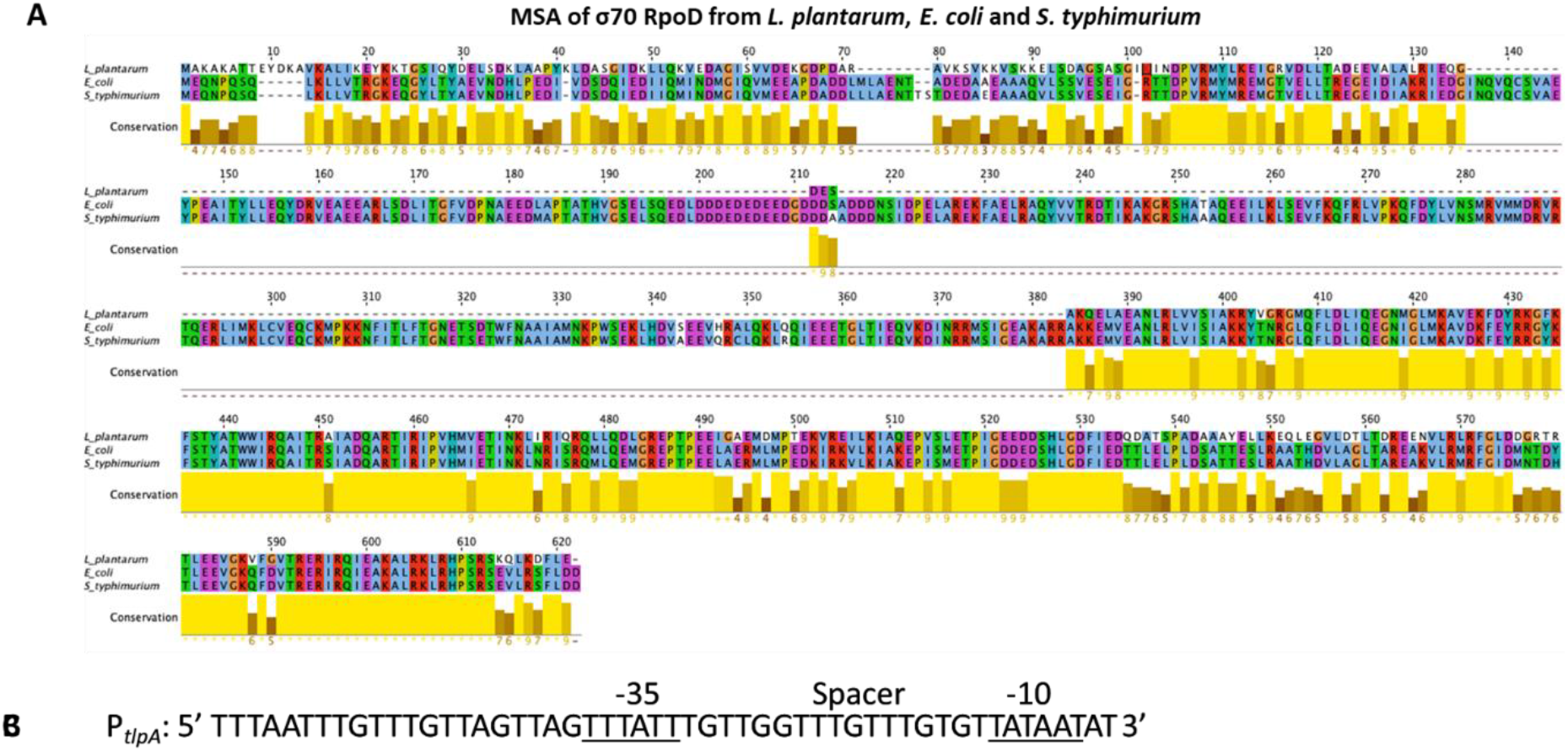
(A) Homology analysis of σ70 RpoD genes from *L. plantarum*, *E. coli*, and *S. typhirmurium*. Height and brightness of the yellow bars indicate the extent to which individual residues are conserved across all 3 bacteria. (B) P_*tlpA*_ promoter sequence with −35, spacer and −10 regions labelled

To understand this, we investigated aspects of *P_tlpA_*’s sequence (Figure 2B). One unique characteristic is that it harbors the sigma70 consensus sequence at the −10 region (5′-TATAAT-3′) but not at the −35 region (5′-TTGACA-3′) (Todt et al., 2012). Based on previous reports in gram positive bacteria like *B. subtilis* (Helmann et al., 1995), it is possible that the deviation from the consensus –35 sequence can be compensated for the presence of the conserved dinucleotide “TG” sequence at the −14 and −15 position of the P_*tlpA*_ promoter. It has been shown that the presence of this sequence upstream of the –10 region in the promoter can mediate rapid promoter melting during transcription initiation and upregulate the transcription rate of corresponding genes. However, most of the promoters reported by Rud et al. (Rud et al., 2006) for *L. plantarum*, also have the conserved “TG” dinucleotide at the –15 position of the promoter. When the strength of the strongest promoter in that library (P_*48*_) was compared to the P_*tlpA*_ promoter, the mCherry production rate by the P_*tlpA*_ promoter was significantly higher. This suggests that the P_*tlpA*_ promoter must have additional reasons that contribute to its exceptional performance in *L. plantarum* WCFS1.

More interestingly, the whole promoter sequence contains no cytosine (C) bases, in contrast to previously reported in *L. plantarum* promoters, most of which contain 2 to 4 cytosine bases in the −35 to −10 region (Rud *et al*., 2006; Meng *et al*., 2021; Sørvig *et al*., 2003; Spangler *et al*., 2019). Additionally, the spacer between the −35 and −10 regions of the P_*tlpA*_ promoter contains no adenine (A) bases. Notably, A and C bases are susceptible to methylation in bacteria, which has been associated with epigenetic gene regulation (Beaulaurier *et al*., 2019; Casadesús *et al*., 2006). However, on analysis of 34 constitutive promoter sequences from the synthetic promoter library reported by Rud *et al*., (2006) and those tested in this study (Supplementary Table S3), no correlation could be derived between promoter strengths and number of C bases within the −35 to −10 region (Supplementary Figure S6A) or the A bases in the spacer (Supplementary Figure S6B). If methylation could be influencing promoter strengths, it would be necessary to identify the methyltransferase recognition sequences in *L. plantarum* to derive meaningful correlations. We then searched for DNA sequences similar to P_*tlpA*_ within the genome of *L. plantarum* WCFS1. Out of 6 hits (Supplementary Figure S7A), only one of them was located upstream of a gene that encodes for a known protein (HAMP domain-containing histidine kinase - locus: lp_0282, complement: 255805..257181), with a percent identity score of 82.76 compared to P_*tlpA*_. This sequence (G**TTTATG**TTGGTTATTTACGTAA**TAAAAT**) was identified as a promoter (referred to as P_*HAMP*_) using BPROM, with −35 and −10 regions (in bold) diverging from P_*tlpA*_ by single bases each (Supplementary Figure S7B). Notably, P_*HAMP*_ also contains four A bases and one C base in the spacer. When the full promoter sequence (Supplementary Table S1) was cloned upstream of mCherry, only weak expression was observed (Supplementary Figure S7C), suggesting that one or more of these mismatches compared to P_*tlpA*_ are essential for driving high-level gene expression. These unique features of the P_*tlpA*_ promoter sequence provide interesting clues for understanding factors affecting promoter strengths in *L. plantarum*. To gain deeper insights into P_*tlpA*_’s unprecedented strength, further studies analyzing mutant libraries of the promoter and/or measuring DNA methylation patterns are required.

### Toxin/Antitoxin based plasmid retention and transient GEMs

Apart from high expression levels, use of lactobacilli for healthcare applications requires strategies to retain heterologous genes in the engineered bacteria in a cheap and compatible manner. TA systems ensure plasmid retention in a bacterial population through a post-segregation killing mechanism. They constitutively express long-lasting toxins and short-lived antitoxins. As long as the plasmid is present, sufficient antitoxin is produced to neutralize the corresponding toxin. On bacterial division, if a daughter cell does not receive any plasmid copies, the antitoxin rapidly degrades, and the active toxin kills the cell. While TA systems have been investigated in the past for bioremediation and biotechnology purposes, their applicability was limited by the fact that their plasmid retention efficiency did not match that of antibiotic or auxotrophy based retention systems (Stirling et al., 2020). However, interest in TA systems has reemerged for living therapeutic applications because of 2 reasons – (i) better understanding of TA systems leading to improved efficiencies (Fedorec *et al*., 2019) and (ii) biosafety features they offer in reducing horizontal gene transfer (Wright *et al*., 2013). Accordingly, reports have recently emerged where TA systems are showing greater promise for bacteria engineered as live vaccines or drug delivery vehicles (Kan et al., 2020; Abedi *et al*., 2022). While these demonstrations have been done in *E. coli*, the use of TA system in lactobacilli for plasmid retention has not yet been systematically investigated. From literature reports and using the TA finder bioinformatics tool, we identified and selected 5 different type II TA system (all named as toxin/antitoxin) – (i) Txe/Axe, from *Enterococcus faecium* that was shown to ensure long-term plasmid retention in *E. coli* (Fedorec *et al*., 2019), (ii) YafQ/DinJ from *L. casei* (Levante *et al*., 2019), (iii) HigB/HigA and (iv) MazF/MazE from *L. plantarum* WCFS1, and (v) HicA/HicB from *L. acidophilus* (Phylogeny in Figure 1A). In all these systems, the toxin is an endoribonuclease and the antitoxin is its corresponding inhibitory protein. These modules were added to the plasmid encoding P_*tlpA*_-driven mCherry expression (Figure 3A) and the resultant strain was repeatedly sub-cultured for up to 100 generations. Plasmid retention was quantified by determining the proportion of the bacterial population expressing mCherry using flow cytometry and agar plate colony imaging analysis (Supplementary Figure S5B). Notably, the sensitivity of this analysis was greatly improved by the high-level of expression driven by the P_*tlpA*_ promoter, which enabled clear demarcation of plasmid-retained and plasmid-lost cells (Figure 3B). Such a clear demarcation was not possible with the other promoters, like P_*23*_ since the fluorescent signal seemed to partially overlap with background signal from non-fluorescent cells (Supplementary Figure S5A). In the absence of a TA system (P_*tlpA*_ mCherry plasmid), the proportion of plasmid-bearing bacteria steadily declined by about 1%/ generation, ending with ~15% of the population retaining the plasmid after 100 generations (Figure 3C). Compared to this, the Txe/Axe system initially supported better retention with a plasmid loss of about 0.5%/generation for 40 generations, after which this loss accelerated to ~1.2%/generation, ending in ~18% of the population retaining the plasmid after 100 generations. HigB/HigA and MazF/MazE systems performed similarly for the most part but provided slightly better retention after 100 generations (20% and 30% respectively). HicA/HicB slowed plasmid loss to 0.5%/generation for 50 generations and 0.8%/ generation, thereafter, resulting in retention level of ~35% after 100 generations. Finally, YafQ/DinJ was found to provide the best retention capabilities with plasmid loss of 0.5%/generation for 70 generations and 1%/ generation thereafter, resulting in a retention level of ~40% after 100 generations (Figure 3C).

**Figure 3.**
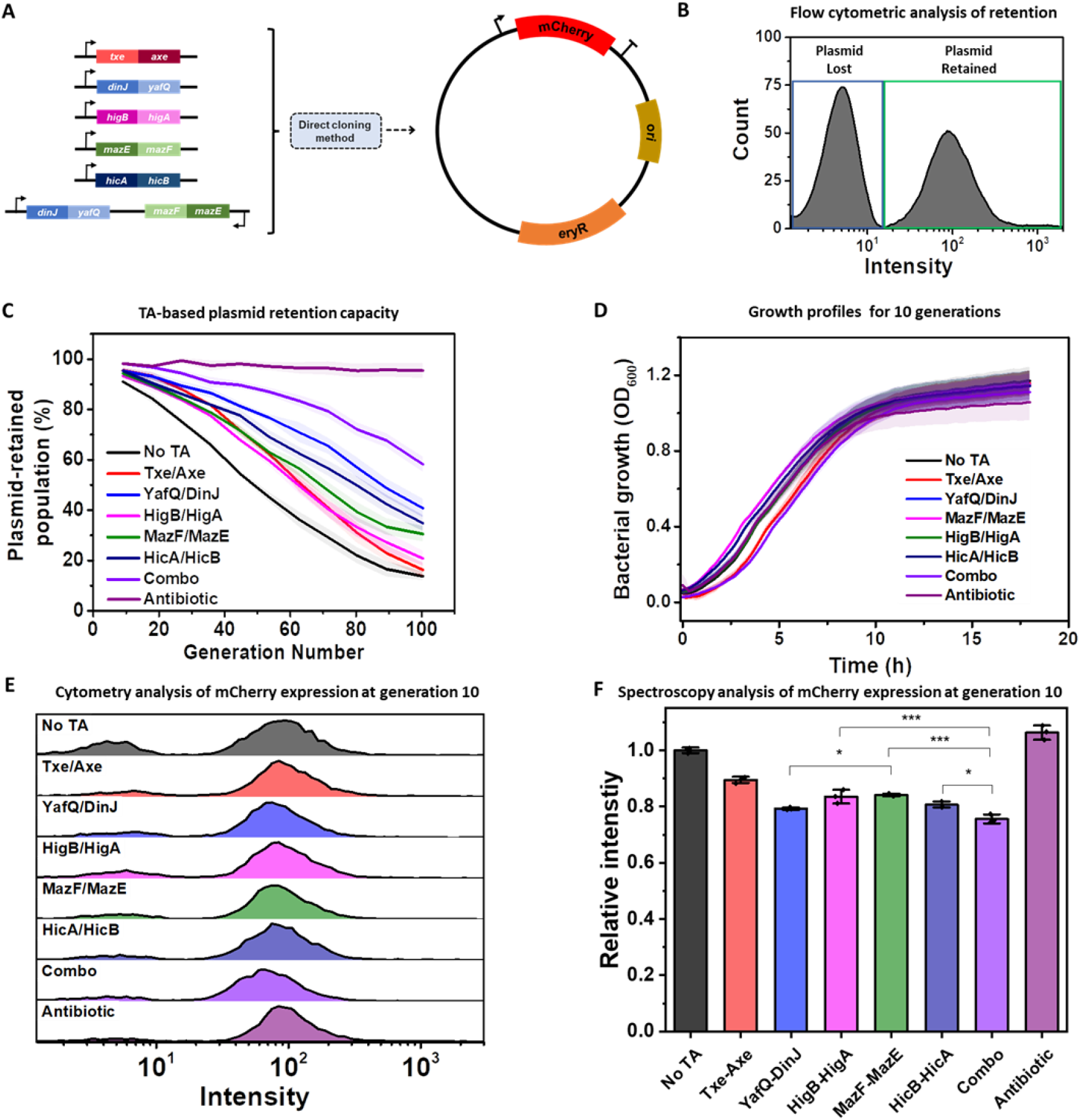
(A) Schematic Representation of cloning the different TA genetic modules into the P_*tlpA*_-mCherry plasmid. (B) Sample flow Cytometry histogram plot of the P_*tlpA*_-mCherry plasmid containing strain without any TA module or selection pressure after 50 generations of serial passaging in the absence of antibiotic. The green box corresponds to the bacterial population retaining the plasmid and the blue box represents the population devoid of the plasmid. (C) Plasmid retention analysis of the TA module containing strains for 100 generations without antibiotics along with no TA and antibiotic selection pressure conditions for comparison. (D) Growth rate (OD_600_) of strains with the TA modules, no TA and antibiotic retention over 10 generations at 37°C. In (C) and (D), the solid lines represent mean values and the lighter bands represents SD calculated from three independent biological replicates. Combo = MazF/MazE + YafQ/DinJ. (E) Flow cytometry plots of strains containing TA modules, no TA and antibiotic retention after 10 generations. The Y-axis for each plot represents counts with plot heights in the range of 450 – 500 (F) Fluorescence Spectroscopy analysis of strains containing TA modules, no TA and antibiotic retention after 10 generations. The relative intensity has been plotted for all the TA strains by normalizing their respective fluorescence values against the “No TA” strain. The data represents three independent biological replicates. p-values are calculated using one-way ANOVA with Tukey test on respective means (* p<0.05, *** p<0.001). The “No TA”, “Txe-Axe” and “Antibiotic” conditions are significantly different from other candidates, so their p - values have not been explicitly highlighted.

Previous studies have shown that combining different TA systems can cumulatively offer better plasmid retention capabilities (Torres et al., 2003; Bardaji *et al*., 2019), although this has not been tested in lactobacilli. So, we combined the best-performing TA system endogenous to *L. plantarum* WCFS1 (MazF/MazE) with the best-performing non-endogenous system (YafQ/DinJ) and observed better plasmid retention capabilities with this combination, yielding a slow plasmid loss of 0.2%/generation for 50 generations and a gradual increase to 0.8%/generation thereafter, resulting in a considerably higher retention of 60% over 100 generations. Comparatively, plasmids maintained under antibiotic selection pressure were steadily retained at >90% through 100 generations, as expected. In all strains harboring TA modules, bacterial growth rates (Figure 3D) and mCherry expression levels (Figure 3E) were found to be minimally impacted compared to “No TA” or antibiotic-retention conditions over the first 10 generations. These results suggest that the toxins did not drastically impede the regular functioning of the cells. Fluorescence spectroscopy analysis of the liquid cultures after 10 generations (Figure 3F) reveals that the TA modules showing higher efficiency in retaining plasmids in the absence of selection pressure (YafQ/DinJ and combo), have significantly lower intensities of mCherry production in comparison to the other TA candidates. The greatest drop in protein expression (~23%) was observed in the strain harboring the TA combo and could be due to an increase in the plasmid size possibly burdening the cells and maybe even resulting in a minor drop in copy number. However, since The YafQ/DinJ construct also causes a drop of comparable magnitude (~20%), it is possible that the toxin in this system mildly interferes with protein expression, which becomes detectable with the overexpression of mCherry by P_*tlpA*_ but does not drastically affect growth. Further in depth investigation would be required to identify the specific cause of this effect. However, it must be noted that even with the drop in expression level caused by the combo TA system, P_*tlpA*_ driven mCherry expression was at least 4-fold higher than that of the next strongest promoter, P_*23*_.

It is important to note that a single generation corresponds to a bacterial duplication, so 10 generations = 2^10^ or ~10^3^ bacteria and 100 generations = 2^100^ or ~10^30^ bacteria from a single cell. Potential applications of lactobacilli for living therapeutics or engineered living materials are not expected to reach such high generation numbers either due to short application time periods (Janahi *et al*., 2018; LeCureux and Dean, 2018; Wang *et al*., 2020) or external growth restrictions (Bhusari *et al*., 2022). Thus, the >90% retention levels provided by the combo TA system for up to 40 generations should be more than sufficient for these applications. Furthermore, loss of the plasmid only reverts the bacteria to their non-GEM probiotic status, thus enabling the generation of transient GEMs that would be desirable for such applications. Accordingly, by varying the TA system used, the GEM lifetime of these organisms could be tuned. Based on this concept, we introduce a new metric, G_50_, for characterizing such transient GEMs. The G_50_ value corresponds to the generation at which half the population of a strain has lost its plasmid. As shown in Figure 4, G_50_ can be tuned from 50 generations for the No TA condition up to 110 generations (extrapolated) for the combo system. Further exploration of additional TA systems in future studies will contribute to more fine tuning of retention lifetimes and possibly even lead to near-perfect retention as has been achieved in *E. coli* by the Txe/Axe system (Fedorec *et al*., 2019). These G_50_ values are expected to depend on culture parameters and environmental factors, due to which it could also become a useful metric for assessing natural and industrial conditions in which lactobacilli grow and function.

**Figure 4.**
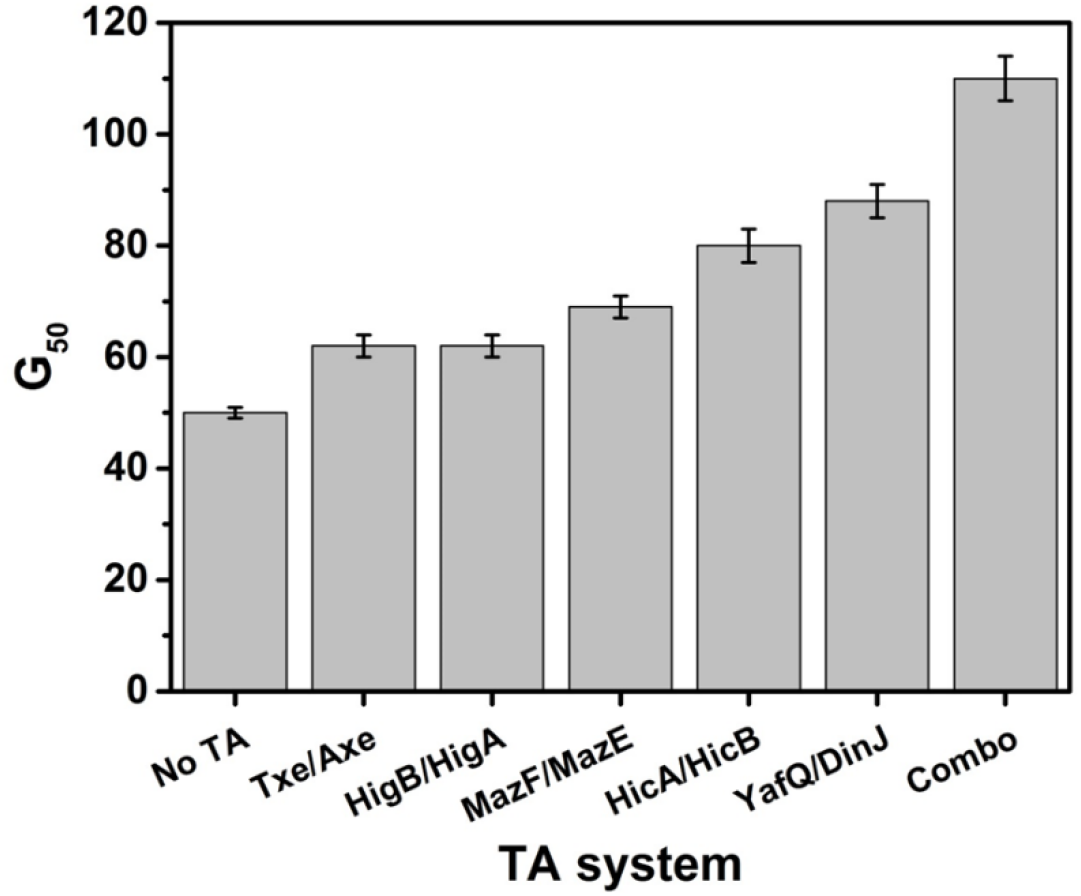
G_50_ values of the different TA systems tested in *L. plantarum*. Combo = MazF/MazE + YafQ/DinJ

## CONCLUSIONS

Lactobacilli as probiotics and commensals in humans and animals have immense potential to be developed for healthcare applications but as non-model organisms, have very poorly equipped genetic toolboxes. Addressing this limitation, this study describes two new genetic modules, characterized in probiotic *L. plantarum* – an ultra-strong constitutive promoter (P_*tlpA*_) and TA plasmid retention systems. Our results demonstrate that the promoter drives gene expression at levels over 5-fold higher than the strongest promoters previously reported in *L. plantarum* and the TA systems decelerate plasmid loss in a tunable manner without the need for external selection pressures or genomic manipulations.

Apart from the impact, these modules will have in expanding the programmability of lactobacilli, the unique conceptual insights gained from this work will aid in the further development of genetic parts. For one, the unique features of the P_*tlpA*_ promoter sequence that originate from phylogenetically distant Salmonella provide clues to understanding what drives promoter strength. Secondly, both homologous and heterologous toxin/antitoxin systems can be used in *L. plantarum* for plasmid retention without considerably affecting bacterial growth rates or protein production levels. More interestingly, the plasmid retention efficacy of these systems can be improved by combining two toxin-antitoxin systems, a phenomenon that has yet been tested only in *E. coli*. Finally, these systems provide the possibility to generate tunable transient GEMs since plasmid loss reverts the cells to their non-GEM probiotic status, characterized by the new G_50_ metric.

## DATA AVAILABILITY

All data are available from the corresponding authors upon reasonable request.

## SUPPLEMENTARY DATA

Supplementary Data are available online.

## ACKNOWLEDGEMENT

The *L. plantarum* WCFS1 strain was a kind gift from Prof. Gregor Fuhrmann (Helmholtz Institute for Pharmaceutical Research, Saarland). The plasmids pSIP403 and pLp_3050sNuc were a kind gift from Prof. Lars Axelsson (Addgene plasmid # 122028) and Prof. Geir Mathiesen (Addgene plasmid # 122030) respectively. The plasmid pTlpA39-Wasabi was a kind gift from Prof. Mikhail Shapiro (Addgene plasmid # 86116). The plasmid pUC-GFP-AT was a kind gift from Prof. Chris Barnes (Addgene plasmid # 133306). The authors thank Dr. Samuel Pearson at the INM – Leibniz Institute for New Materials for discussions that led to the G_50_ concept.

## REFERENCES

Abedi, M.H., Yao, M.S., Mittelstein, D.R., Bar-Zion, A., Swift, M.B., Lee-Gosselin, A., et al. (2022) Ultrasound-controllable engineered bacteria for cancer immunotherapy. Nat. Commun., 13, 1–11.

Allison, G.E. and Klaenhammer, T.R. (1996) Functional analysis of the gene encoding immunity to lactacin F, lafI, and its use as a Lactobacillus-specific, food-grade genetic marker. Appl. Environ. Microbiol., 62, 4450–4460.

Ashraf, R. and Shah, N.P. (2011) Selective and differential enumerations of Lactobacillus delbrueckii subsp. bulgaricus, Streptococcus thermophilus, Lactobacillus acidophilus, Lactobacillus casei and Bifidobacterium spp. in yoghurt--A review. Int. J. Food Microbiol., 149, 194–208.

Bardaji, L., Añorga, M., Echeverría, M., Ramos, C. and Murillo, J. (2019) The toxic guardians—multiple toxin-antitoxin systems provide stability, avoid deletions and maintain virulence genes of Pseudomonas syringae virulence plasmids. Mobile DNA., 10, 1–17.

Beaulaurier, J., Schadt, E.E. and Fang, G. (2019) Deciphering bacterial epigenomes using modern sequencing technologies. Nat. Rev. Genet., 20,157–172.

Bhusari, S., Sankaran, S. and Del Campo, A. (2022) Regulating bacterial behavior within hydrogels of tunable viscoelasticity. Adv. Sci., 9, p2106026.

Bibalan, M.H., Eshaghi, M., Rohani, M., Esghaei, M., Darban-Sarokhalil, D., Pourshafie, M.R., et al. (2017) Isolates of Lactobacillus plantarum and L. reuteri display greater antiproliferative and antipathogenic activity than other Lactobacillus isolates. J. Med. Microbiol., 66, 1416–1420.

Bron, P.A., Hoffer, S.M., Van Swam, I.I., De Vos, W.M. and Kleerebezem, M. (2004) Selection and characterization of conditionally active promoters in Lactobacillus plantarum, using alanine racemase as a promoter probe. Appl. Environ. Microbiol., 70, 310–317.

Casadesús, J. and Low, D. (2006) Epigenetic gene regulation in the bacterial world. Microbiol. Mol. Biol. Rev., 70, 830–856.

Castillo-Hair, S.M., Baerman, E.A., Fujita, M., Igoshin, O.A. and Tabor, J.J. (2019) Optogenetic control of Bacillus subtilis gene expression. Nat. Commun., 10, 1–11.

Chan, M.Z.A., Chua, J.Y., Toh, M. and Liu, S.Q. (2019) Survival of probiotic strain Lactobacillus paracasei L26 during co-fermentation with S. cerevisiae for the development of a novel beer beverage. Food Microbiol., 82, 541–550.

Chee, W.K.D., Yeoh, J.W., Dao, V.L. and Poh, C.L. (2022) Highly reversible tunable thermal-repressible split-t7 RNA polymerases (thermal-T7RNAPs) for dynamic gene regulation. ACS Synthetic Biology, 11(2), pp.921–937.

Chen, Y., Qi, M., Xu, M., Huan, H., Shao, W. and Yang, Y. (2018) Food-grade gene transformation system constructed in Lactobacillus plantarum using a GlmS-encoding selection marker. FEMS Microbiol. Lett., 365, fny254.

Courbet, A., Endy, D., Renard, E., Molina, F. and Bonnet, J. (2015) Detection of pathological biomarkers in human clinical samples via amplifying genetic switches and logic gates. Sci. Transl. Med., 7, 289ra83–289ra83.

Darby, T.M., and Jones, R.M. (2017) Beneficial influences of Lactobacillus plantarum on human health and disease. In Floch, M., Ringel, Y., Walker,W.A. (ed.), The microbiota in gastrointestinal pathophysiology. Academic Press, Vol. I, pp. 109–117.

Davis, M.C., Kesthely, C.A., Franklin, E.A. and MacLellan, S.R. (2017) The essential activities of the bacterial sigma factor. Can. J. Microbiol., 63, 89–99.

Dawoud, T.M., Davis, M.L., Park, S.H., Kim, S.A., Kwon, Y.M., Jarvis, N., O’Bryan, C.A., Shi, Z., Crandall, P.G. and Ricke, S.C. (2017) The potential link between thermal resistance and virulence in Salmonella: a review. Frontiers in veterinary science, 4, p.93.

de Vos, W.M. (2011) Systems solutions by lactic acid bacteria: from paradigms to practice. Microb. Cell Fact., 10, 1–13.

du Toit, M., Engelbrecht, L., Lerm, E. and Krieger-Weber, S. (2011) Lactobacillus: the next generation of malolactic fermentation starter cultures—an overview. Food Bioprocess Technol., 4, 876–906.

Edgar, R.C. (2004) MUSCLE: a multiple sequence alignment method with reduced time and space complexity. BMC Bioinf., 5, 1–19.

Elowitz, M.B. and Leibler, S. (2000) A synthetic oscillatory network of transcriptional regulators. Nature., 403, 335–338.

Fedorec, A.J., Ozdemir, T., Doshi, A., Ho, Y.K., Rosa, L., Rutter, J., et al. (2019) Two new plasmid post-segregational killing mechanisms for the implementation of synthetic gene networks in Escherichia coli. iscience., 14, 323–334.

Gaida, S.M., Sandoval, N.R., Nicolaou, S.A., Chen, Y., Venkataramanan, K.P. and Papoutsakis, E.T. (2015) Expression of heterologous sigma factors enables functional screening of metagenomic and heterologous genomic libraries. Nat. Commun., 6, 1–10.

Grady, R. and Hayes, F. (2003) Axe–Txe, a broad-spectrum proteic toxin–antitoxin system specified by a multidrug-resistant, clinical isolate of Enterococcus faecium. Mol. Microbiol., 47, 1419–1432.

Halbmayr, E., Mathiesen, G., Nguyen, T.H., Maischberger, T., Peterbauer, C.K., Eijsink, V.G., et al. (2008) High-level expression of recombinant β-galactosidases in Lactobacillus plantarum and Lactobacillus sakei using a sakacin P-based expression system. J. Agric. Food Chem., 56, 4710–4719.

Heiss, S., Hörmann, A., Tauer, C., Sonnleitner, M., Egger, E., Grabherr, R. et al. (2016) Evaluation of novel inducible promoter/repressor systems for recombinant protein expression in Lactobacillus plantarum. Microb. Cell Factories., 15, 1–17.

Helmann, J.D. (1995) Compilation and analysus of Bacillus Subtilis σ A-dependent promoter sequences: evidence for extended contact between RNA polymerse and upstream promoter DNA. Nucleic Acids Res. 23, 2351–2360.

Hurme, R., Berndt, K.D., Normark, S.J. and Rhen, M. (1997) A proteinaceous gene regulatory thermometer in Salmonella. Cell., 90, 55–64.

Janahi, E.M.A., Haque, S., Akhter, N., Wahid, M., Jawed, A., Mandal, R.K., et al. (2018) Bioengineered intravaginal isolate of Lactobacillus plantarum expresses algal lectin scytovirin demonstrating anti-HIV-1 activity. Microb. Pathog., 122, 1–6.

Jiménez, E., Fernández, L., Maldonado, A., Martín, R., Olivares, M., Xaus, J., et al. (2008) Oral administration of Lactobacillus strains isolated from breast milk as an alternative for the treatment of infectious mastitis during lactation. Appl. Environ. Microbiol., 74, 4650–4655

Kasımoğlu, A., Göncüoğlu, M. and Akgün, S. (2004) Probiotic white cheese with Lactobacillus acidophilus. Int. Dairy J., 14, 1067–1073.

Kan, A., Gelfat, I., Emani, S., Praveschotinunt, P. and Joshi, N.S. (2020) Plasmid vectors for in vivo selection-free use with the probiotic E. coli Nissle 1917. ACS synthetic biology, 10(1), pp.94–106.

Landete, J.M., Langa, S., Revilla, C., Margolles, A., Medina, M. and Arqués, J.L. (2015) Use of anaerobic green fluorescent protein versus green fluorescent protein as reporter in lactic acid bacteria. Appl. Microbiol. Biotechnol., 99, 6865–6877.

LeCureux, J.S. and Dean, G.A. (2018) Lactobacillus mucosal vaccine vectors: immune responses against bacterial and viral antigens. mSphere., 3, e00061–18.

Letunic, I. and Bork, P. (2007) Interactive Tree Of Life (iTOL): an online tool for phylogenetic tree display and annotation. Bioinformatics., 23, 127–128.

Levante, A., Folli, C., Montanini, B., Ferrari, A., Neviani, E. and Lazzi, C. (2019) Expression of DinJ-YafQ System of Lactobacillus casei group strains in response to food processing stresses. Microorganisms., 7, 438.

Ma, B., Forney, L.J. and Ravel, J. (2012) The vaginal microbiome: rethinking health and diseases. Annu. Rev. Microbiol., 66, 371.

Mastromarino, P., Macchia, S., Meggiorini, L., Trinchieri, V., Mosca, L., Perluigi, M., et al. (2009) Effectiveness of Lactobacillus-containing vaginal tablets in the treatment of symptomatic bacterial vaginosis. Clin. Microbiol. Infect., 15, 67–74.

Mathiesen, G., Øverland, L., Kuczkowska, K. and Eijsink, V.G. (2020) Anchoring of heterologous proteins in multiple Lactobacillus species using anchors derived from Lactobacillus plantarum. Sci. Rep., 10, 1–10.

Mathiesen, G., Sveen, A., Brurberg, M.B., Fredriksen, L., Axelsson, L. and Eijsink, V.G. (2009) Genome-wide analysis of signal peptide functionality in Lactobacillus plantarum WCFS1. BMC genomics., 10, 1–13.

Madeira, F., Pearce, M., Tivey, A., Basutkar, P., Lee, J., Edbali, O., et al. (2022) Search and sequence analysis tools services from EMBL-EBI in 2022. Nucleic Acids Res.

Meier-Kolthoff, J.P. and Göker, M. (2019) TYGS is an automated high-throughput platform for state-of-the-art genome-based taxonomy. Nat. Commun., 10, 1–10.

Meng, Q., Yuan, Y., Li, Y., Wu, S., Shi, K. and Liu, S. (2021) Optimization of electrotransformation parameters and engineered promoters for Lactobacillus plantarum from wine. ACS Synth. Biol., 10, 1728–1738.

Nguyen, T.T., Mathiesen, G., Fredriksen, L., Kittl, R., Nguyen, T.H., Eijsink, V.G., et al. (2011) A food-grade system for inducible gene expression in Lactobacillus plantarum using an alanine racemase-encoding selection marker. J. Agric. Food Chem., 59, 5617–5624.

Nguyen, H.M., Pham, M.L., Stelzer, E.M., Plattner, E., Grabherr, R., Mathiesen, G., Peterbauer, C.K., Haltrich, D. and Nguyen, T.H. (2019) Constitutive expression and cell-surface display of a bacterial β-mannanase in Lactobacillus plantarum Microb. Cell Factories, 18,1–12.

Paget, M.S. and Helmann, J.D. (2003) The σ70family of sigma factors. Genome Biol., 4, 1–6.

Pedrolli, D.B., Ribeiro, N.V., Squizato, P.N., de Jesus, V.N., Cozetto, D.A., Tuma, R.B., et al. (2019) Engineering microbial living therapeutics: the synthetic biology toolbox. Trends Biotechnol., 37, 100–115.

Piraner, D.I., Abedi, M.H., Moser, B.A., Lee-Gosselin, A. and Shapiro, M.G. (2017) Tunable thermal bioswitches for in vivo control of microbial therapeutics. Nat. Chem. Biol., 13, 75–80.

Plessas, S., Fisher, A., Koureta, K., Psarianos, C., Nigam, P. and Koutinas, A.A. (2008) Application of Kluyveromyces marxianus, Lactobacillus delbrueckii ssp. bulgaricus and L. helveticus for sourdough bread making. Food Chem., 106, 985–990.

Rosenfeldt, V., Benfeldt, E., Nielsen, S.D., Michaelsen, K.F., Jeppesen, D.L., Valerius, N.H., et al. (2003) Effect of probiotic Lactobacillus strains in children with atopic dermatitis. J. Allergy Clin. Immunol., 111, 389–395.

Rottinghaus, A.G., Ferreiro, A., Fishbein, S.R., Dantas, G. and Moon, T.S. (2022) Genetically stable CRISPR-based kill switches for engineered microbes. Nature communications, 13(1), pp.1–17.

Rud, I., Jensen, P.R., Naterstad, K. and Axelsson, L. (2006) A synthetic promoter library for constitutive gene expression in Lactobacillus plantarum. Microbiology., 152, 1011–1019.

Russo, P., Iturria, I., Mohedano, M.L., Caggianiello, G., Rainieri, S., Fiocco, D., et al. (2015) Zebrafish gut colonization by mCherry-labelled lactic acid bacteria. Appl. Microbiol. Biotechnol., 99, 3479–3490.

Salomé-Desnoulez, S., Poiret, S., Foligné, B., Muharram, G., Peucelle, V., Lafont, F., et al. (2021) Persistence and dynamics of fluorescent Lactobacillus plantarum in the healthy versus inflamed gut. Gut Microbes., 13, 1897374.

Siezen, R.J. and van Hylckama Vlieg, J.E. (2011) Genomic diversity and versatility of Lactobacillus plantarum, a natural metabolic engineer. Microb. Cell Factories., 10, 1–13.

Stirling, F. and Silver, P.A.(2020) Controlling the implementation of transgenic microbes: are we ready for what synthetic biology has to offer? Molecular cell, 78(4), pp.614–623.

Sørvig, E., Grönqvist, S., Naterstad, K., Mathiesen, G., Eijsink, V.G. and Axelsson, L. (2003) Construction of vectors for inducible gene expression in Lactobacillus sakei and L. plantarum. FEMS Microbiol. Lett., 229, 119–126.

Sørvig, E., Mathiesen, G., Naterstad, K., Eijsink, V.G. and Axelsson, L. (2005a) High-level, inducible gene expression in Lactobacillus sakei and Lactobacillus plantarum using versatile expression vectors. Microbiology., 151, 2439–2449.

Sørvig, E., Skaugen, M., Naterstad, K., Eijsink, V.G. and Axelsson, L. (2005b) Plasmid p256 from Lactobacillus plantarum represents a new type of replicon in lactic acid bacteria, and contains a toxin–antitoxin-like plasmid maintenance system. Microbiology, 151, 421–431.

Spangler, J.R., Caruana, J.C., Phillips, D.A. and Walper, S.A. (2019) Broad range shuttle vector construction and promoter evaluation for the use of Lactobacillus plantarum WCFS1 as a microbial engineering platform. Synth. Biol., 4, ysz012.

Spath, K., Heinl, S. and Grabherr, R. (2012) Direct cloning in Lactobacillus plantarum: electroporation with non-methylated plasmid DNA enhances transformation efficiency and makes shuttle vectors obsolete. Microb. Cell Factories., 11, 1–8.

Takala, T. and Saris, P. (2002) A food-grade cloning vector for lactic acid bacteria based on the nisin immunity gene nisI. Appl. Microbiol. Biotechnol., 59, 467–471.

Tauer, C., Heinl, S., Egger, E., Heiss, S. and Grabherr, R. (2014) Tuning constitutive recombinant gene expression in Lactobacillus plantarum. Microb. Cell Factories., 13, 1–11.

Teughels, W., Durukan, A., Ozcelik, O., Pauwels, M., Quirynen, M. and Haytac, M.C. (2013) Clinical and microbiological effects of Lactobacillus reuteri probiotics in the treatment of chronic periodontitis: a randomized placebo-controlled study. J. Clin. Periodontol., 40, 1025–1035.

Todt, T.J., Wels, M., Bongers, R.S., Siezen, R.S., Van Hijum, S.A. and Kleerebezem, M. (2012) Genome-wide prediction and validation of sigma70 promoters in Lactobacillus plantarum WCFS1. PLoS One., 7, e45097

Tran, A.M., Unban, K., Kanpiengjai, A., Khanongnuch, C., Mathiesen, G., Haltrich, D. and Nguyen, T.H. (2021) Efficient secretion and recombinant production of a lactobacillal α-amylase in Lactiplantibacillus plantarum WCFS1: analysis and comparison of the secretion using different signal peptides. Front. Microbiol., 12, 689413.

Turroni, F., Ventura, M., Buttó, L.F., Duranti, S., O’Toole, P.W., Motherway, M.O.C., et al. (2014) Molecular dialogue between the human gut microbiota and the host: a Lactobacillus and Bifidobacterium perspective. Cell. Mol. Life Sci., 71,183–203.

Van Kranenburg, R., Golic, N., Bongers, R., Leer, R.J., De Vos, W.M., Siezen, R.J., et al. (2005) Functional analysis of three plasmids from Lactobacillus plantarum. Appl. Environ. Microbiol., 71, 1223–1230.

Wang, M., Fu, T., Hao, J., Li, L., Tian, M., Jin, N., et al. (2020) A recombinant Lactobacillus plantarum strain expressing the spike protein of SARS-CoV-2. Int. J. Biol. Macromol., 160, 736–740.

Wang, B., Kitney, R.I., Joly, N. and Buck, M. (2011) Engineering modular and orthogonal genetic logic gates for robust digital-like synthetic biology. Nat. Commun., 2, 1–9.

Waterhouse, A.M., Procter, J.B., Martin, D.M., Clamp, M. and Barton, G.J. (2009) Jalview Version 2—a multiple sequence alignment editor and analysis workbench. Bioinformatics., 25, 1189–1191.

Watterlot, L., Rochat, T., Sokol, H., Cherbuy, C., Bouloufa, I., Lefèvre, F., et al. (2010) Intragastric administration of a superoxide dismutase-producing recombinant Lactobacillus casei BL23 strain attenuates DSS colitis in mice. Int. J. Food Microbiol., 144, 35–41.

Wright, O., Stan, G.B. and Ellis, T.(2013) Building-in biosafety for synthetic biology. Microbiology, 159(Pt_7), pp.1221–1235.

Xie, Y., Wei, Y., Shen, Y., Li, X., Zhou, H., Tai, C et al. (2018) TADB 2.0: an updated database of bacterial type II toxin–antitoxin loci. Nucleic Acids Res., 46, D749–D753.

Yamaguchi, Y. and Inouye, M. (2011) Regulation of growth and death in Escherichia coli by toxin–antitoxin systems. Nat. Rev. Microbiol., 9, 779–790.

Zheng, J., Wittouck, S., Salvetti, E., Franz, C.M., Harris, H.M., Mattarelli, P., et al. (2020) A taxonomic note on the genus Lactobacillus: Description of 23 novel genera, emended description of the genus Lactobacillus Beijerinck 1901, and union of Lactobacillaceae and Leuconostocaceae. Int. J. Syst. Evol. Microbiol., 70, 2782–2858.

Zocco, M. A., Verme L. Z.D., Cremonini, F., Piscaglia, A.C., Nista, E.C., Candelli, M., et al. (2006) Efficacy of Lactobacillus GG in maintaining remission of ulcerative colitis. Aliment. Pharmacol. Ther., 23, 1567–1574.

