## Supporting information for "Novel genetic modules encoding high-level antibiotic-free protein expression in probiotic lactobacilli"

### Plasmid Construction Scheme

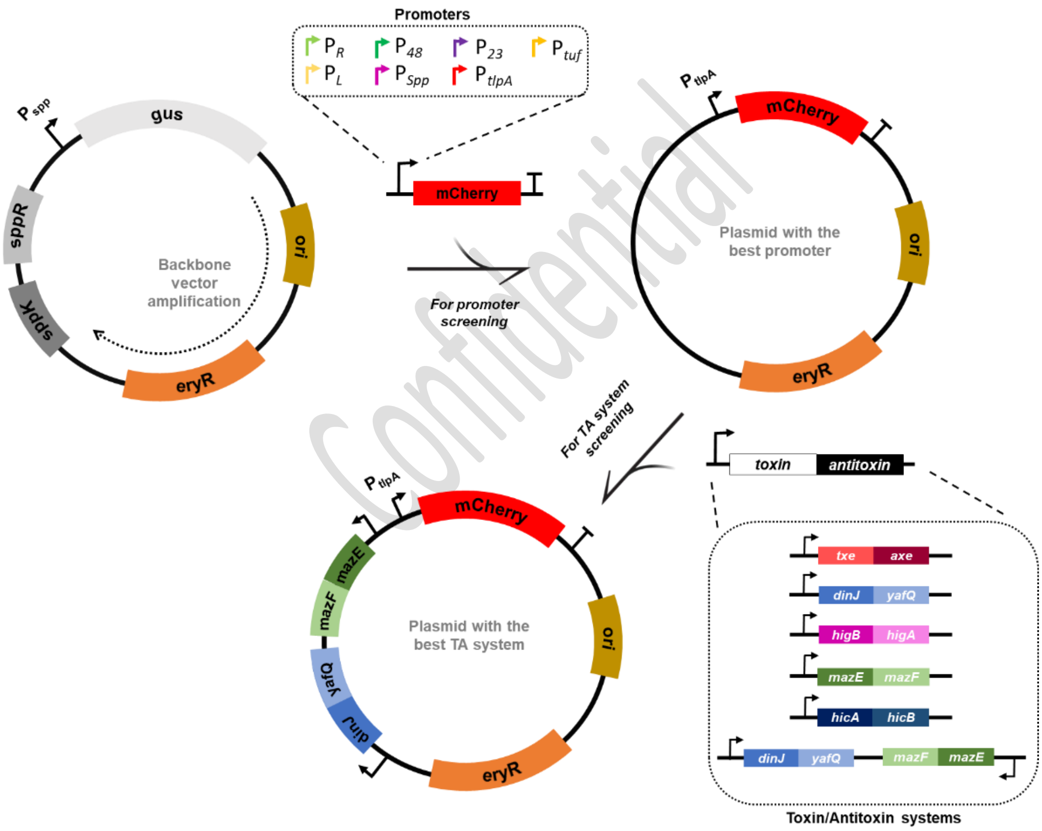

Confidential

**Supplementary Figure S1** Construction of plasmid variants using the direct cloning approach developed in this study. The p256 origin of replication and the erythromycin resistance marker has been amplified from the pSIP403 vector backbone. The interchangeable promoter sequences were upstream of the mCherry reporter gene with $P_{tlpA}$ promotter showing the best performance (highlighted). The activity of the toxin/antitoxin modules were checked in the pTlpA mcherry plasmid with the combo TA system showing the best plasmid retention (higlighted).

**Supplementary Table S1.** Nucleotide Sequences of the common genetic parts and interchangeable promoter sequences tested in this study

| NAME | GENETIC PART | COMMON FEATURE | ORIENTATION | SEQUENCE |
|---|---|---|---|---|
| Ori p256 | Origin of replication | Yes | → | ttgagatcctttttttctgcgcgtaatctgctgcttgcaa<br>acaaaaaaaccaccgctaccagcggtggtttgtttgcc<br>ggatcaagagctaccaactctttttccgaaggtaactg<br>gcttcagcagagcgcagataccaaatactgttcttcta<br>gtgtagccgtagttaggccaccacttcaagaactctgt<br>agcaccgcctacatacctcgctctgctaatcctgttacc<br>agtggctgctgccagtggcgataagtcgtgtcttaccg<br>ggttggactcaagacgatagttaccggataaggcgca<br>gcggtcgggctgaacggggggttcgtgcacacagccc<br>agcttggagcgaacgacctacaccgaactgagatacc<br>tacagcgtgagctatgagaaagcgccacgcttcccgaa<br>gggagaaaggcggacaggtatccggtaagcggcagg<br>gtcggaacaggagagcgcacgagggagcttccaggg<br>ggaaacgcctggtatctttatagtcctgtcgggtttcgc<br>cacctctgacttgagcgtcgatttttgtgatgctcgtca<br>ggggggcggagcctatggaaa |
| eryR | Antibiotic resistance | Yes | → | ttgaacaaaaatataaaatattctcaaaactttttaac<br>gagtgaaaaagtactcaaccaaataataaaacaatt<br>gaatttaaaagaaaccgataccgtttacgaaattgga<br>acaggtaaagggcatttaacgacgaaactggctaaa<br>ataagtaaacaggtaacgtctattgaattagacagtca<br>tctattcaacttatcgtcagaaaaattaaaactgaata<br>ctcgtgtcactttaattcaccaagatattctacagtttca<br>attccctaacaaacagaggtataaaattgttgggaata<br>ttccttacaatttaagcacacaaattattaaaaaagtg<br>gtttttgaaagccgtgcgtctgacatctatctgactgttg<br>aagaaggattctacaagcgtaccttggatattcaccga<br>acactagggttgctcttgcacactcaagtctcgattcag<br>caattgcttaagctgccagcggaatgctttcatcctaa<br>accaaaagtaaacagtgtcttaataaaacttacccgc<br>cataccacagatgttccagataaatattggaagctata<br>taagtactttgtttcaaaatgggtcaatcgagaatatcg<br>tcaactgtttactaaaaatcagtttcgtcaagcaatga<br>aacacgccaaagtaaacaatttaagtaccattacttat<br>gagcaagtattgtctatttttaatagttatctattattta<br>acgggaggaaataa |
| mCherry | Reporter gene | No | → | atggtttcaaagggtgaagaagataacatggctatca<br>tcaaggaattcatgcgtttcaaggttcacatggaaggt<br>tcagttaacggtcacgaattcgaaatcgaaggtgaag<br>gtgaaggtcgtccatacgaaggtactcaaactgctaa<br>gttaaaggttactaagggtggtccattaccattcgcttg<br>ggatatcttatcaccacaattcatgtacggttcaaaggc<br>ttacgttaagcacccagctgatatcccagattacttaaa |

Confidential

| | | | | |
|---|---|---|---|---|
| | | | | gttatcattcccagaaggtttcaagtgggaacgtgttatgaacttcgaagatggtggtgttgttactgttactcaagattcatcattacaagatggtgaattcatctacaaggttaagttacgtggtactaacttcccatcagatggtccagttatgcaaaagaagactatgggttgggaagcttcatcagaacgtatgtacccagaagatggtgctttaaagggtgaaatcaagcaacgtttaaagttaaaggatggtggtcactacgatgctgaagttaagactacttacaaggctaagaagccagttcaattaccaggtgcttacaacgttaacatcaagttagatatcacttcacacaacgaagattacactatcgttgaacaatacgaacgtgctgaaggtcgtcactcaactggtggtatggatgaattatacaagtaa |
| RBS | Ribosome-binding site | No | → | tttgtttaactttaagaaggaga |
| $P_{tlpA}$ | Promoter | No | → | tttaatttgtttgttagttagtttatttgttggtttgtttgtgttataatat |
| $P_{spp}$ | Promoter | No | → | gcccatattaacgtttaaccgataaagttgaacgttaatatttttttt |
| $P_{48}$ | Promoter | No | → | tcgtaagttgttgacatggaacgaggaatgtgataatctgtgagt |
| $P_{23}$ | Promoter | No | → | ctgatgacaaaaagagaaaattttgataaaatagtcttagaattaaattaaaaa |
| $P_{Tuf}$ | Promoter | No | → | tctgtttacaaatcagattaggctatatataatatttaagga |
| $P_{L}$ | Promoter | No | → | ttgacataaataccactggcggtgatact |
| $P_{R}$ | Promoter | No | → | ttgactattttacctctggcggtgataa |
| P*HAMP* | Promoter | No | → | caaaatgtcgttgacgtttatgttggttatttacgtaataaaatcacgac |
| T7 ter | Terminator | No | → | ctagcataaccccttggggcctctaaacgggtcttgaggggttttttg |

Confidential

**Supplementary Table S2.** Nucleotide Sequences of the toxin/antitoxin modules tested in this study

| NAME | GENETIC PART | COMMON FEATURE | ORIENTATION | SEQUENCE |
|---|---|---|---|---|
| YafQ/DinJ | Toxin-Antitoxin operon | No | → | agatggcagttacgcttccatttgtgcgaagaaaaagcca<br>gaaatccggttgaattcatctcacatgatgttatcccactgt<br>gtttacattgggataattcgtgatataattaggttgttagat<br>aggaaggagtggttagcaatggcagccacaaagaaaga<br>aactcgcttgaatattcgtgttgatccggaattaaaaagt<br>gctgctcaaatcgtagcaaatgatatgggcatcgacttga<br>ccgcagctgttactatgttcatgacgaaaatggtgaaaga<br>tcacgccctcccgtttaccccaacaagtctaccagttgaaa<br>ccttacaggcgctgaaagaagcaaagcacccagagctgc<br>tcaaaaaatacagcacgcctgatgacatgtggagagact<br>tgaatgtatagtctggttccgacgcctacatttaagcgcga<br>tctaaaacgactctccaagaagcattggccgatggacga<br>actaaagacggctgttaatctcctagctgctggtacaaat<br>gctgaactattaagcaaaaagtatgcagatcatgccttgt<br>cttcaagcagcgagtggaaaggatatcgtgaactacatg<br>ttgacggccctcgtggcgactggttgctaatctataaaatt<br>aagcagcaagatctcattttgaccctggttag<br>aactggatctcatcataaccttttgggtaaatagaactacg<br>aaaggccgtccaaaaaaggcggcttttgtgttgtggtcaa<br>ccagcagattcattcgtcaatatacacgtgacttaacgcca<br>atcttcaatcagagattagaaaccgcgaactcgtcggccc<br>agtaaaaatgttcagctatcgagatgagttgggtgatctg<br>gggaagtgagcatcaataagatgccaatctttatcatgca<br>gggttggaacgaaaatttaacccaatg |
| MazF/MazE | Toxin-Antitoxin operon | No | ← | gaaggatttacggtcaacgagtgtctttcggacacgtccgt<br>tgatattgttatttgactcgtggtcttactccaatcaattat<br>actgacaagcatcgcaaatataaaaccgccctacctaac<br>ctgagtaagttacagggcgaagtagtgctgattagaatgg<br>aaagtcgaagttgtaatacaccgtttgcgcgacgtgataa<br>aaatcggcactgcgcatggtttcgatgtatgtgatactccg<br>tcctgctctgggggtcgcatcaaacgaataaatttgggca<br>gcagcaatttgtccgtgggtattgtaaccattaagactaa<br>agtagccgggcaaatcacgcattgtagacgtaattggtg<br>aaacaattacaaacccggtgttctgagtgtagagattact<br>actaagaactacagctggccgacgtttctttatttcgcggc<br>cacgttgtgggtcaaaatcaatccagatgatgtctttttgt<br>ttaggtaaataagtcatttactcgacctcattccagtcgac<br>atcggaccatgcttcagtcattggccgtttcggcttggagc<br>cggcataaggattctcgatacgtggtacgagtgtatatga<br>ctcaccatcacttgatgggatgagtaaccatcgttgtcctt<br>tcttaaaacgaccgtctttaggtagacttaatgccaatga<br>attaccacgactaactgtctttacctcaatcatatcaaggc<br>ctcctcgtattaataaacacactatttatgcaacgagtat<br>aacacgtattacgaaataagacagctaaagcaatcgcg |

Confidential

| | | | | |
|---|---|---|---|---|
| | | | | ataactaattgat<br>aagcacacggtttttagtaattcatcaataaaggaagag<br>actaattaatgtcgccgtaactcagtgttatcaagttggat<br>ggcccgctttagtattttccacctttagcctctt |
| HicA/HicB | Toxin-Antitoxin operon | No | → | tactgatgatgaacatgccccagcctgttgtagctctttgc<br>gcaataaatcatttattttgtctggattgaatagagcttga<br>gcaaaatctttggtaaaatcattcatgaaggagttcttctt<br>tctgtatgtgtttttttgttcaaaccaatcatacgagaaag<br>gaactcctttttctaatgtttaaagaaataaatttaagga<br>atcattcatccttacacaaactattttacagtctcattctt<br>cataaaccacatcctagaaaagaactattaccatatcaa<br>atcaaacaaattcaagataaaatcaaagagatgaata<br>aagatgaaaaaaactaatcaatatatgaagtataagg<br>gttatgaaggttcaattgaatatactttggaggataaga<br>ttctttttggtaaggttcaggggattaagagtcttatttctt<br>acgaaggtaatacaatagatgaattggagaaggattttc<br>aaggagccattgatgattatctaatgagctgtaaggaag<br>atggagttataccagagaaacctttaaaggtaattttaa<br>tgtgcgcattgatccttcattacatgaaaaattggctaat<br>tatgctgcaacgaagcaccaatcattgaatgctagtgta<br>gaagaagctataaaaagatttttggcttaggtgaaaga<br>cgcttggaatttttcaagcgtttttttcatgaaaacaattg<br>ctaaattcattgaatgagtgtactattttagtataacaat<br>agttgaacgggtgatgtcggcgatatagg |
| HigB/HigA | Toxin-Antitoxin operon | No | → | tactgatgatgaacatgcccactctggttggttcctagcta<br>aaataggggtcgtactagtcgggtaatcagatgaattgg<br>atatttgggtgtggtcagctaataataggagcaagagtc<br>ggatggattgagatatacacgtgatacgggtataatcga<br>ttaagaattttgcagataaggaaactcacaaggtttacc<br>aacaaaaattctctaaacaattgccaccaaccattcagc<br>aattagcattgcgaaaattactaatgattgatcatgcgga<br>aacaattaatgatttgagcctaccgcctgccaatcacttg<br>gaaaaattaagtcatgatcgccaagggcaatatagcatc<br>aggattaataatcagtatcggatatgttttgcaattcgaa<br>atggtaatgagttttatgatgttgaaatagtcgattatcat<br>catggttaggagaccaaaaaatgaatgaaatcccaaca<br>cctaaaattagtgaaattcttgaagaagagtttatggctc<br>ccttgcatatctctgcatattttttggctcaacagattgggg<br>tacccacgtcacggattcaagacttactacatgatcgtcga<br>caggtgacggttgatacatcgctgcggctaggacgattctt<br>tggagtgtcggatcgttatttcttggaacttcaaaatgata<br>ttgaaattcgtaatttgaaacagatacatggtgctgaata<br>tgcacagataaaaaagtatcaagtcagttaaataagga<br>cgtagaccgtagttgcttggtagcgtactacggtttttgca<br>aatataacctaaactgataagcagcagatcttaaatttta<br>aggttatcaacaggtggataaagcagtggcgtttgtatta<br>tccacctgttgataatcattgagttggttaattgtactttca<br>ctactgggttaactggtcgtgagaaaaatggtcaagcatt<br>agtttcgtggtgatgtcggcgatatagg |

Confidential

| | | | | |
|---|---|---|---|---|
| Txe/Axe | Toxin-Antitoxin operon | No | ← | acgcgtaacaaacacattcattaataacaatcaacacta<br>agctatattagctttttaaatgactagggtaaataaaata<br>aggatcaaaaacttttaagtttctgaccctttccttacttcc<br>gattggttaatagtgatcttttgcagaataaataaatatc<br>gtttcattttcaactctatatatcagtctatgttcatctgta<br>attcttctggaccattttccagataaatcatgctttaatgg<br>ctcaggttttcctaatccagcaaagggggaacgatcgat<br>atcttttattaacttgttaatctttttatattgcttttgtttc<br>cttgctcatgccaataaagataatcatcccaagcatcatc<br>agaccaagccttaatcatcagattcaacctcgattaagt<br>catgtgttttaaatgcacctttggagaattgttcatctcctc<br>gacgaatttttccatgacgtaattattagaaagtgttctc<br>aacgtttcttgcatagaatcataatctctttttgataatac<br>aacaactgtatcttctacatctttacttgttacaataagtg<br>tttcagcatcctcattaacttgtttcatataactacgtaaa<br>ttttggcggaaatttgaataagctactgcttccattccttt<br>cacccttatttatttctataaaacaattgtacattatatt<br>gtacaattaagcaaatgattaatattctaacttcttataa<br>ttaaaagtgctatcacttttagtttttaggataaaaggtg<br>acgtcactttttgagtgctggtacctcaaatcctgctttt |

Confidential

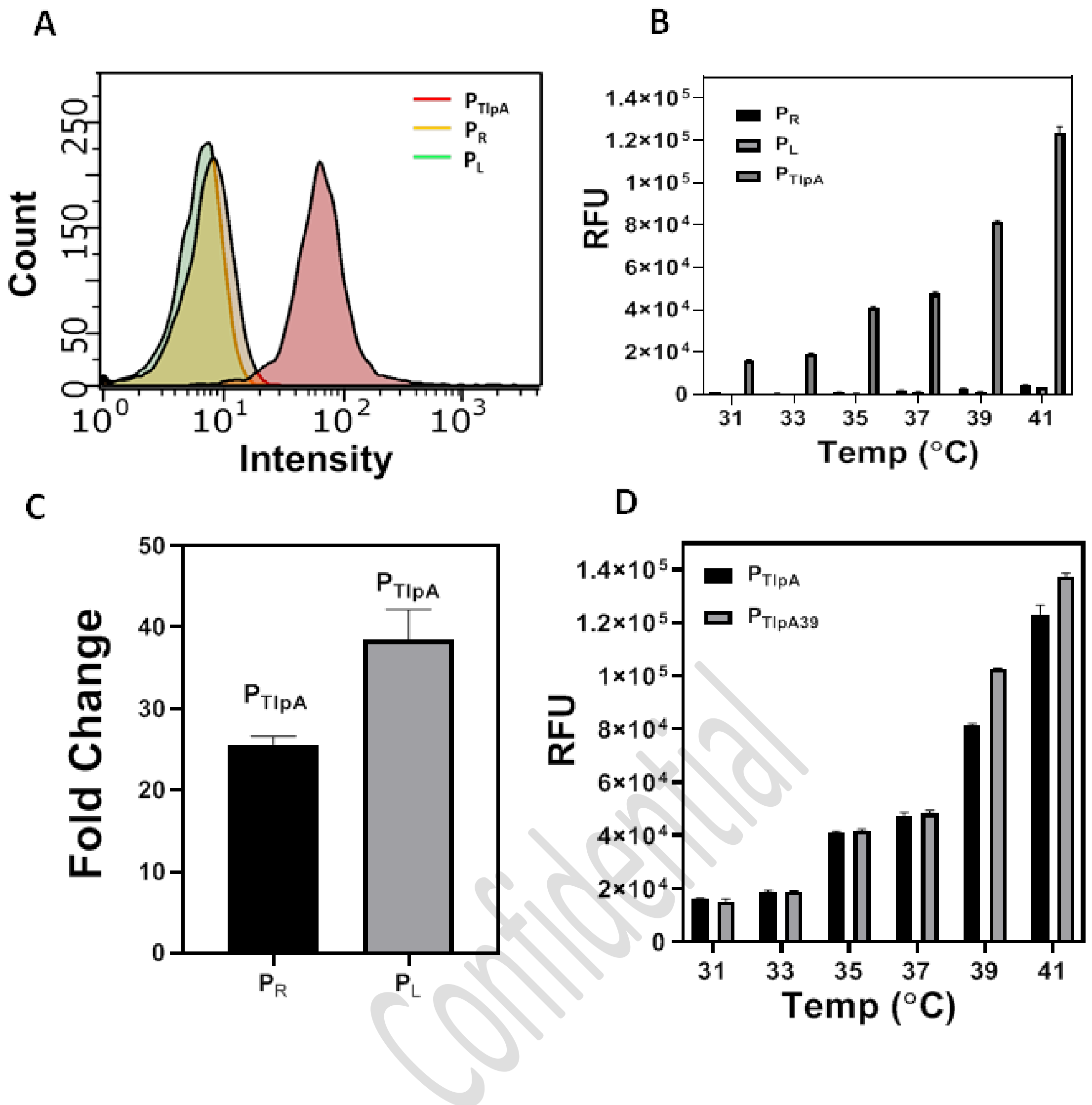

Confidential

**Supplementary Figure S2.** (A) Flow Cytometry analysis of $P_R$, $P_L$ and $P_{tlpA}$ driven mCherry expression in *L. plantarum* WCFS1 after 18 h incubation at 37°C. (B) Fluorescence Spectroscopy analysis of the $P_R$, $P_L$ and $P_{tlpA}$ driven mCherry expression after 18 h incubation in the Thermocycler setup at thermal gradients ranging from 31°C to 41°C (See Materials and Methods section "Microplate reader Setup for Thermal Gradient Analysis"**)**. (C) Fold Change of $P_{tlpA}$ driven mCherry expression in comparison to $P_R$ and $P_L$ promoters at 37°C (D) RFU Plot of mcherry production by plasmids pTlpA and pTlpA39 after 18 h incubation at thermal gradients ranging from 31°C to 41°C. The pTlpA39 plasmid encodes for an additional $P_{48}$ promoter-driven codon-optimized TlpA39 repressor. The data in B, C and D corresponds to three independently conducted biological replicates with column heights representing means and whiskers representing SD.

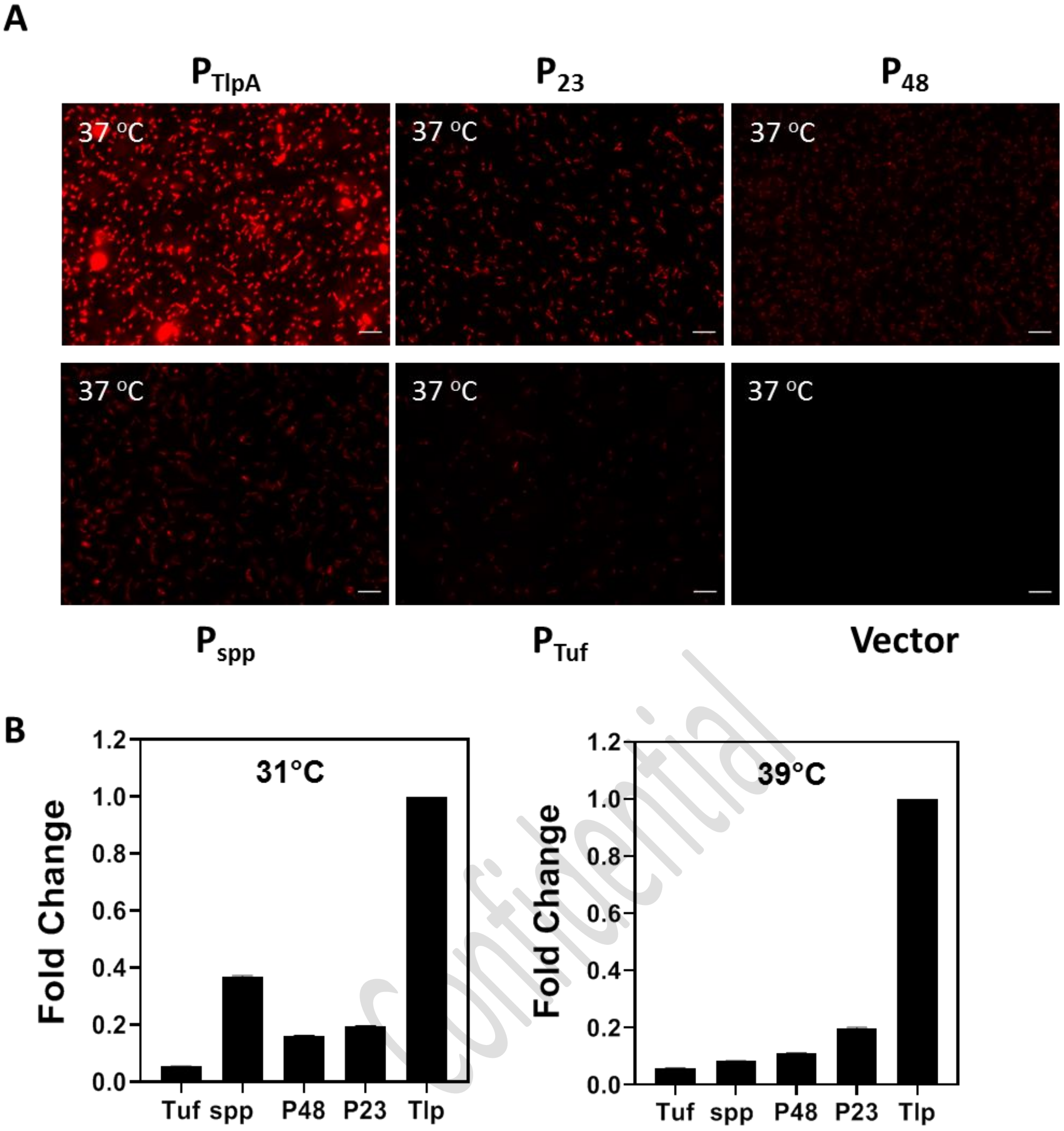

Confidential

**Supplementary Figure S3.** (A) Fluorescence microscopy images of $P_{tlpA}$, $P_{23}$, $P_{48}$, $P_{spp}$, $P_{tuf}$ driven mCherry expression and Empty Vector constructs in *L. plantarum* WCFS1 cultivated at 37°C for 18 h (Scale = 10µm). (B) Fold Change of the $P_{tuf}$, $P_{23}$, $P_{48}$, $P_{spp}$, driven mCherry expression normalized to the expression level of the $P_{tlpA}$ promoter at 31°C and 39°C respectively. The data corresponds to three independently conducted biological replicates with whiskers as SD.

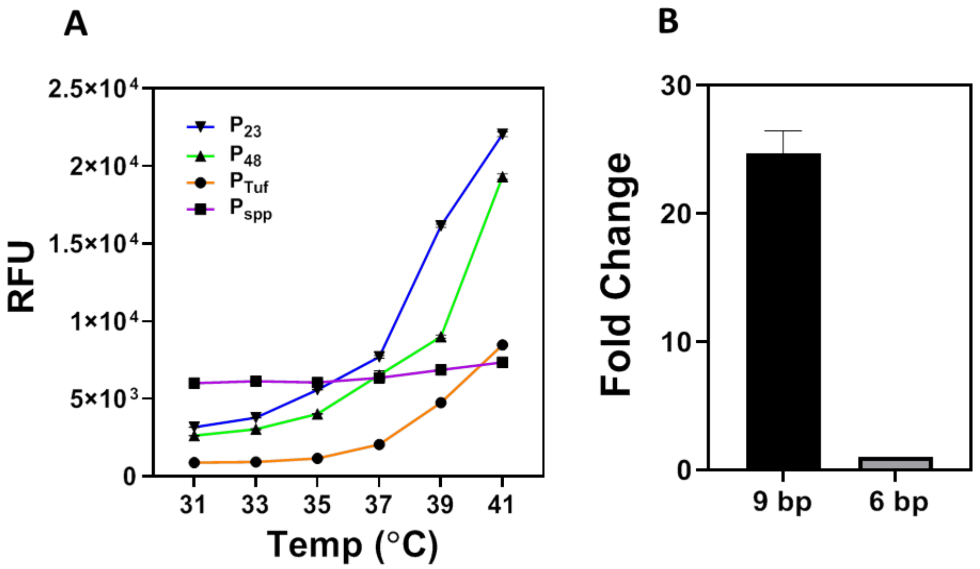

**Supplementary Figure S4.** (A) RFU Plot of $P_{23}$, $P_{48}$ and $P_{tuf}$ driven mCherry expression showing enhanced fluorescence expression in contrast to $P_{spp}$ within the thermal gradients ranging from 31°C to 41°C. The data corresponds to three independently conducted biological replicates with whiskers as SD. (B) Fold Change of the $P_{tlpA}$ promoter driven mCherry expression with 9 bp spacer (between the RBS and the start codon) in comparison to the $P_{tlpA}$ -6 bp spacer construct.

Confidential

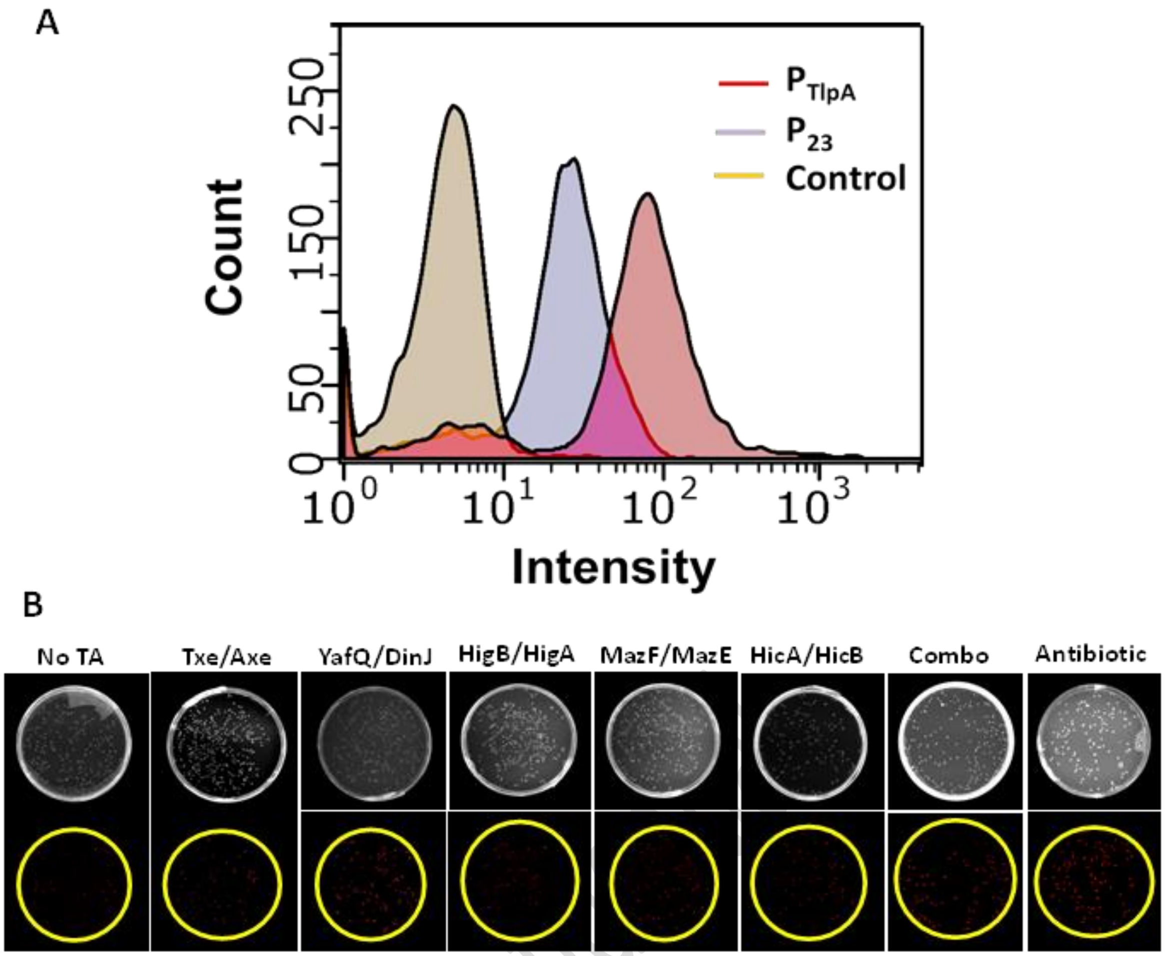

**Supplementary Figure S5.** (A) Flow cytometry plots showing that strains with $P_{23}$-driven mCherry expression produces relatively low intensities with part of the population overlapping with the signal gained from bacteria that do not express any fluorescent proteins (control). In comparison, the signal from $P_{tlpA}$ is clearly demarcated from that of control. (B) Agar-plate based analysis of plasmid retention provided by the different TA systems. The total number of colonies can be determined from the brightfield images and the red colonies visible in the fluorescent images represent the plasmid-retaining bacteria.

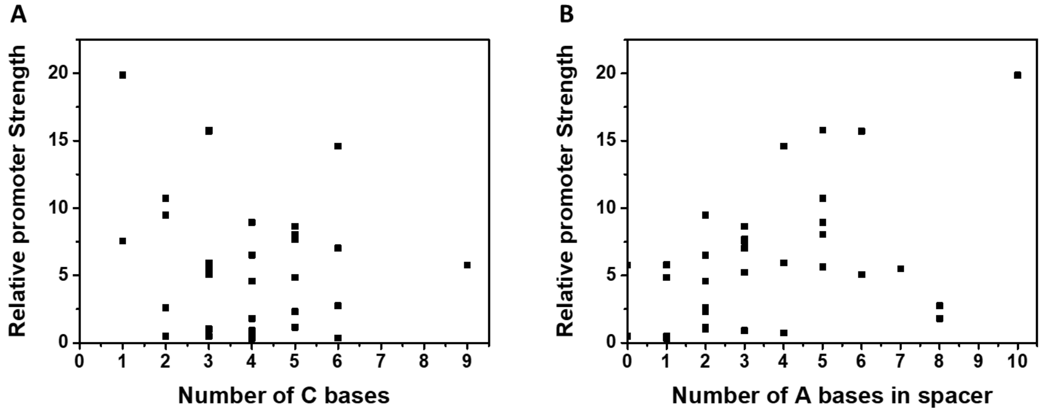

**Supplementary Figure S6.** Plots representing relative promoter strengths of 34 promoters in Table S3 vs (A) number of C bases in the -35 to -10 region and (B)number of A bases in the spacer between the -35 and -10 region (non-bold region in Table S3).

**Supplementary Table S3:** Promoter sequences included in Figure S7 and their relative strengths. Sequences and strengths derived from reference "Rud, I., et al., *A synthetic promoter library for constitutive gene expression in Lactobacillus plantarum.* Microbiology, 2006. **152**(4): p. 1011-1019." The relative strengths of $P_{23}$ and $P_{Tuf}$ from this study were estimated based on expression levels we determined in relation to $P_{48}$.

| Promoter | Sequence | Relative strength |
|---|---|---|
| $P_{20}$ | **-35 TTGACA**ACCTGTGGGCGGTTTGA**TTTGTT -10** | 0.33 |
| $P_{17}$ | **-35 TTGACA**CTGATCCCGGCTGGTGG**TAAATT -10** | 0.35 |
| $P_{13}$ | **-35 TTGACA**GCGTGGGTTGGTGCTGG**TAATTT -10** | 0.48 |
| $P_{41}$ | **-35** -**TGACA**GGGCTGTGATGGTGTGG**TATTGT -10** | 0.50 |
| $P_{43}$ | **-35 TTGACA**GGATAAAGGTCGCCTGG**TATGGT -10** | 0.74 |
| $P_{40}$ | **-35 TCAACA**TACATGGATCG---TGG**TATGTT -10** | 0.91 |
| $P_{44}$ | **-35 TTGACA**CTTGGAGGGTTTGATGG**TAATCT -10** | 1.02 |
| $P_{10}$ | **-35 TTGACA**TTGTTAATGGCCCCTGA**TATATT -10** | 1.15 |
| $P_{34}$ | **-35 TTGACA**GACAAACATAAGGATGA**TATGCT -10** | 1.80 |
| $P_{9}$ | **-35 TTGACA**GGGCTGAGCTGACCTGG**TATGGT -10** | 2.33 |
| $P_{42}$ | **-35 TTGACA** GAGTTGTCTGGTATTGA**TATTGT -10** | 2.60 |
| $P_{5}$ | **-35 TTGACA**CACAAAACCAGACATGG**TATTAT -10** | 2.75 |
| $P_{33}$ | **-35 TTGACA**TGTGACCCGTTTTATGG**TATTAT -10** | 4.58 |
| $P_{21}$ | **-35 TTGACA**TTCGGGGCATCTCGTGG**TATAAT -10** | 4.84 |
| $P_{26}$ | **-35 TGGACA**AGTGATAAAACGGGTGA**TATGAT -10** | 5.06 |
| $P_{22}$ | **-35 TTGACA**GATTAGGGCGGTCATGG**TAAAAT -10** | 5.23 |
| $P_{18}$ | **-35 TTGACA**GGGAGGCTTCGTTGTGA**TAAGAT -10** | 5.79 |
| $P_{47}$ | **-35 TTGACA**GACGCGGGAGAATATGG**TAAAGT -10** | 5.65 |
| $P_{16}$ | **-35 TTGACA**TCCCCCTCTCTTTCTGG**TATAAT -10** | 5.76 |
| $P_{3}$ | **-35 TTGACA**AAGAAGCCGGGTTTTGG**TATAAT -10** | 5.93 |
| $P_{29}$ | **-35 TTGACA**GTTCTGGCTGGATATGG**TAAACT -10** | 6.51 |

Confidential

| | | | | |
|---|---|---|---|---|
| $P_{6}$ | -35 | **TTGACA**GGCAGCCATCTCTATGG**TAAAAT** | -10 | 7.04 |
| $P_{38}$ | -35 | **TTGACA**GAATGTTTTTGTAGTGG**TATAAT** | -10 | 7.56 |
| $P_{30}$ | -35 | **TTGACA**GCCGAGGTACCATGTGG**TATAAT** | -10 | 7.69 |
| $P_{31}$ | -35 | **TTGACA**AAAGTCCCAGGGTATGA**TATACT** | -10 | 8.04 |
| $P_{8}$ | -35 | **TTGACA**AATTCCGGTGTCTATGG**TATTCT** | -10 | 8.64 |
| $P_{25}$ | -35 | **TTGACA**TTAAGGCACATCATTGA**TATGGT** | -10 | 8.94 |
| $P_{35}$ | -35 | **TTGACA**TGCAGGGAGTTTGTTGG**TATAAT** | -10 | 9.49 |
| $P_{1}$ | -35 | **TTGACA**GGGATATTAGAGTATGG**TATGCT** | -10 | 10.74 |
| $P_{4}$ | -35 | **TTGACA**CGCGCAGCAAGGCATGA**TATAAT** | -10 | 14.60 |
| $P_{11}$ | -35 | **TTGACA**GAATGGACATACTATGA**TATATT** | -10 | 15.70 |
| $P_{48}$ | -35 | **TTGACA**TGGAACGAGGAATGTGA**TAATCT** | -10 | 15.79 |
| $P_{23}$ | -35 | **ATGACA**AAAAGAGAAAATTTTGA**TAAAAT** | -10 | 19.86 |
| $P_{Tuf}$ | -35 | **TTTACA**AATCAGATTAGGCTATA**TATAAT** | -10 | 5.49 |

**A**

| Hit | Upstream a gene | Position | Sequence | Sequence length | Identity Matrix |
|---|---|---|---|---|---|
| Hit 1 | Yes | 257,256..257,284 | GTTTATGTTGGTTATTTACGTAATAAAAT | 29 | 82.76 |
| Hit 2 | No | 1,292,083..1,292,110 | ATTTTTAAAACAACTAACAAAAATGAAC | 28 | 69.23 |
| Hit 3 | No | 1,541,627..1,541,653 | GTTATTTTTTAGTTTGTTGTGAAATTT | 27 | 77.78 |
| Hit 4 | No | 2,772,508..2,772,536 | ATACTTAAACACAACAAAAAACACTTAAC | 29 | 32.82 |
| Hit 5 | No | 3,129,614..3,129,645 | ATATGGTCAGTCAATGAAATCAACAAATAAAC | 32 | 60.00 |
| Hit 6 | No | 3,209,426..3,209,455 | ATAGTAACCACCAGCACACCACAAGTAAAC | 30 | 71.43 |

**B**

```
PHAMP   GTTTA--TGTTGG-TTATTTACGTAATAAAAT
PTlpA   -TTTATTTGTTGGTTTGTTTGTGTTATAATAT
         ****  ****** ** ***   ** **** **
```

**C**

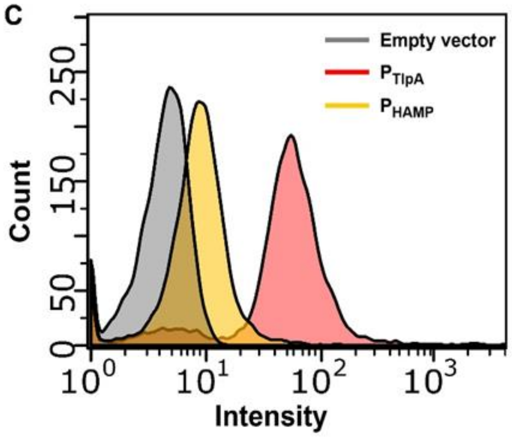

**Supplementary Figure S7.** (A) Results obtained after screening the entire genome of *L. plantarum* WCFS1 for a Pt*lpA*-like sequence. (B) Sequence alignment of $P_{HAMP}$ and $P_{tlpA}$ promoters. (C) Flow Cytometry analysis of $P_{tlpA}$ and $P_{HAMP}$ driven mCherry expression in *L. plantarum* WCFS1 after 18 h incubation at 37°C.
